# Quantification of ligand and mutation-induced bias in EGFR phosphorylation in direct response to ligand binding

**DOI:** 10.1101/2023.02.13.528340

**Authors:** Daniel Wirth, Ece Özdemir, Kalina Hristova

## Abstract

The 58 receptor tyrosine kinases (RTKs) are critically important for human development, and are implicated in many growth disorders and cancers. Here we introduce a methodology to identify and quantify bias in RTK signal transduction across the plasma membrane, and to quantify phosphorylation efficiencies, without contributions from feedback loops and system bias. We show that RTK biased signaling can occur in the first step of signal transduction not only in response to different ligands, but as a consequence of RTK pathogenic mutations as well. Ligand bias and mutation-induced bias are uncoupled here based on a comprehensive data set of dose response curves acquired for three ligands, EGF, TGFα and epiregulin, for wild-type EGFR and for the oncogenic L834R EGFR mutant found in patients with non-small-cell lung cancer (NSCLC). Ligand bias has been extensively studied for GPCRs, and has revolutionized the GPCR field. The demonstration of pathogenic mutation-induced bias in RTK signal transduction across the plasma membrane will open new avenues for the exploration of RTK biased inhibitors as highly specific anti-cancer therapies.

## INTRODUCTION

Receptor tyrosine kinases (RTKs) are single pass membrane receptors which control cell growth, differentiation, motility, survival, and metabolism. They have been implicated in many diseases, and are valuable drug targets. RTKs transduce biochemical signals via lateral interactions in the plasma membrane, by forming catalytically active dimers (1–3). RTK dimerization, which is modulated by ligand binding, brings the kinase domains together in close proximity so they cross-phosphorylate each other on tyrosines in the activation loop (3,4). This activates the kinases and they phosphorylate additional tyrosines which serve as binding sites for effector molecules, thus triggering downstream signaling cascades. Recent work has suggested that the signaling pathways originating from different RTK tyrosines can be differentially activated by different ligands - a phenomenon known as “ligand bias” (4).

Ligand bias has been studied primarily in the context of G-protein coupled receptors (5–7), where it has been shown to originate in the first step of signal transduction across the plasma membrane, i.e., due to differential signal propagation across the length of the receptor (8–10). For RTKs, the origin of ligand bias has been debated (11,12). Furthermore, experiments meant to explore RTK ligand bias have thus far probed for “functional selectivity” (13,14), which is “the combined effect of ligand and system bias” (15). While ligand bias is universal and pertains to all cell types as it depends on receptors and ligands, its manifestation may be different in different cells and tissues due to the system bias (15). System bias is determined by the cellular/tissue/ physiological state context, including the identities of downstream signaling effectors in cells (15). For RTKs, it has been further shown that the abundances of signaling effectors can introduce system biases (16). Importantly, system bias can be perceived even at the level of RTK phosphorylation, which can be affected by feedback loops that operate within the cell (15,17). A recent review of the current state of the field (15) emphasizes that “biased signaling represents very complex pharmacology, making experiment design, interpretation and description challenging and often inconsistent— causing confusion about what has really been measured and what can be concluded”. Here we introduce a methodology to measure ligand bias in direct response to ligand binding, thus simplifying interpretation and gaining insights into the origin of the bias. We show that we can identify and quantify ligand bias without contributions from feedback loops or system bias; we call it “intrinsic ligand bias”. We also show that we can identify and quantify intrinsic signaling bias that is induced by an RTK pathogenic mutation. The new methodology utilizes automatic imaging and data processing, and can be used for high-throughput screening of biased inhibitors to eliminate the deleterious effects of pathogenic RTK mutations.

## RESULTS

### A model system to measure RTK phosphorylation in direct response to ligand binding

To be able to measure RTK phosphorylation in direct response to ligands without contribution from feedback loops and systems bias, we used plasma membrane derived vesicles produced via osmotic vesiculation (18,19). Such vesicles are produced from cells that have been transfected with genes encoding RTKs labeled with fluorescent proteins, and are imaged in a confocal microscope. As described in Supplemental Data, we developed a neural network approach to analyze the vesicle images. Towards this goal, we trained a computer vision Faster R-CNN network with ~1400 labeled vesicles. The program identified the vesicles (Figure S1), and quantifies the fluorescence intensity on the membrane, as well as the fluorescence intensity inside and outside the vesicles.

Previous work has shown that the lipid composition of the vesicles is very similar to the lipid composition of the plasma membrane (19). While all plasma membrane derived vesicles are known to have defects that allow the passage of macromolecules through the membrane (20), the vesicles produced via osmotic vesiculation allow the passage of very large macromolecules (19). As a result, cytoplasmic signaling proteins such as Grb2-GFP (MW = 60 kDa) and PLCγ-GFP (MW = 210 kDa) diffuse through the vesicle membranes and become infinitely diluted in the buffer which is contiguous with the vesicle lumens (19). So do cytoplasmic proteins that bind to the lipids on the cytoplasmic side (such as PLCδ-PH-GFP), or to RTKs on the cytoplasmic side (such as PLCγ-GFP). These proteins dissociate from the membrane because their residence times on the membrane are much shorter than the timescale of vesicle production, ~ 12 hours (19). Thus, these vesicles lack the components of signaling feedback loops.

Here we investigated the permeability of these vesicles to FITC-labeled dextrans (20 – 2000 kDa) which were added externally after vesicle production. The FITC intensity inside and outside the vesicles were measured after 1 hour to quantify the degree of dextran penetration through the vesicle membrane. Figures 1A and 1B shows that the 20 kDa and 70 kDa dextrans equilibrate across the vesicle membrane without any obstruction (intensity ratio: 1.00 ± 0.03 and 0.98 ± 0.03, respectively). This confirms the presence of large defects in the membrane, and explains the lack of retention of cytoplasmic signaling proteins. Reduced penetration of dextrans is observed starting from molecular weight 250 kDa (intensity ratio: 0.84 ± 0.05). Dextrans are known to form rodlike structures in aqueous solutions (21), and hydrodynamic radii for the 20, 73, 250, 500, and 2000 kDa dextrans have been reported as 3.2, 6.5, 11.5, 15.9, and 26.9 nm, respectively (22). Taking into account that the hydrodynamic radius for an IgG antibody (150 kDa) is 5.4 nm, and thus falls in between the hydrodynamic radii of 20 kDa and 70 kDa dextran, we predict that antibodies which are added externally will penetrate the vesicles. This makes it possible to detect specific phosphotyrosines on the kinase domain of an RTK in the vesicle lumen as illustrated in Figure S3. The antibodies are recruited to the vesicle membrane upon tyrosine phosphorylation, where their fluorescence intensities are measured.

**Figure 1.**
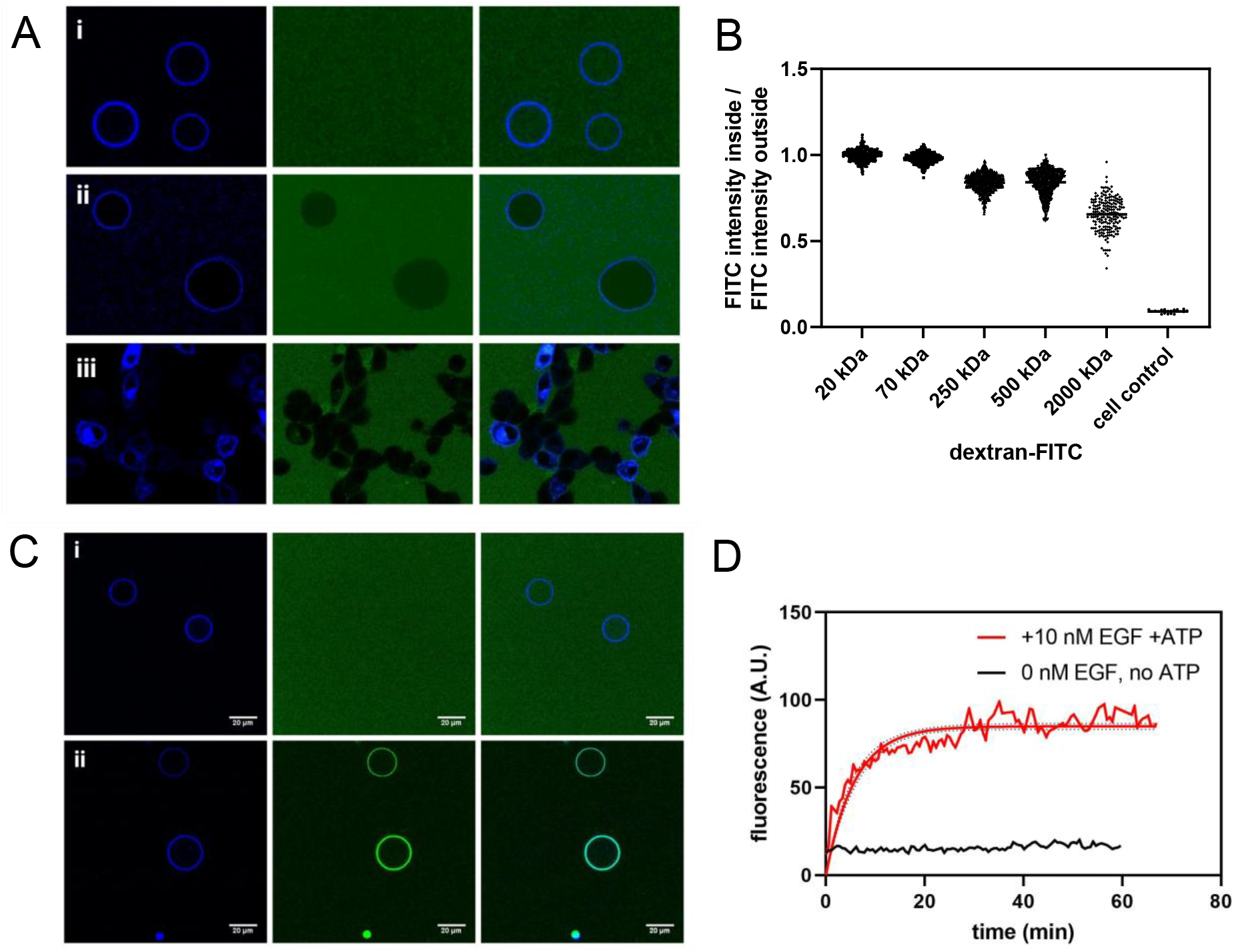
Plasma membrane derived vesicles as a tool to probe RTK phosphorylation in response to ligand binding. **(A)** Determination of the size cutoff for equilibration across the vesicle membrane. FITC-labeled dextrans of different molecular weights were added to vesicles with FGFR3-mTurquoise in their membrane. a) Left column: receptor, middle column: dextran, right column: overlay. i) 70 kDa dextran. The intensity inside the vesicle and outside is the same. ii) 2000 kDa dextran. The intensity inside the vesicle is lower. iii) cells, EGFR-mTurquoise + control antibody (rat IgG2bκ-FITC). The antibody does not cross the cell membrane. **(B)** Intensity ratios between FITC-labeled dextran inside and outside of vesicles for dextrans of different molecular weight. Each data point represents the ratio for one vesicle. Intensities for ~3400 vesicles were measured. **(C)** EGFR phosphorylation in vesicles. First column: receptor channel, middle column: antibody channel, right column: overlay. i) Vesicles derived from CHO cells with EGFR-mTurq incorporated into the vesicle membrane were incubated with 10 μg/mL IgG-FITC isotype control antibody. ii) Vesicles derived from CHO cells with EGFR-mTurq incorporated into the vesicle membrane were incubated with 100 nM EGF, ATP/salt cocktail, and FITC anti-pY 4G10 antibody. The antibody fluorescence can be seen on the membrane. **(D)** Phosphorylation signal on the vesicle membrane over time. The fluorescence of the antibody was measured on the membrane of a single vesicle over time in response to 10 nM EGF with added ATP cocktail (red line). The black line shows a control experiment in the absence of EGF and ATP.

Vesicles were produced from cells transfected with EGFR. Experiments were set up with 100 nM EGF in the presence of an ATP cocktail containing Mg^2+^ and a phosphatase inhibitor (see Materials and Methods). A FITC-labeled anti-pY 4G10 antibody, which recognizes any phosphorylated tyrosine residue on EGFR in response to EGF stimulation, was used for detection. An IgG-FITC isotype control antibody was used as a control. While no significant binding to the vesicle membrane was detected for the control antibody, a clear increase in membrane fluorescence was observed for the anti-pY antibody upon EGF addition (Figure 1C). In both cases the solution fluorescence intensity of the antibody fluorophores is equal inside and outside the vesicles, showing that antibodies are able to freely diffuse across the vesicle membrane.

The maximum antibody binding and therefore receptor phosphorylation is achieved after about 20 minutes, with a time course which is likely affected by ligand and antibody diffusion (Figure 1D). Importantly for this work, a plateau in the fluorescence was observed, demonstrating that the phosphorylation comes to equilibrium, consistent with the expectation that no feedback loops are present. As a control, no increase in fluorescence was observed in the absence of ligand and ATP kinase cocktail.

### Quantification of intrinsic ligand bias in EGFR signal propagation across the plasma membrane

We investigated if there is preference for the phosphorylation of one of two tyrosines when EGFR is activated by three EGFR ligands: EGF, TGFα, and epiregulin. The two tyrosines that were probed, Y1068 and Y1173, are in the long unstructured tail of EGFR and have profound importance for signaling. Phosphorylation of Y1068 leads to the recruitment of Grb2 and Gab1 and the activation of AKT and STAT3/5 signaling pathways (23–25). On the other hand, Y1173 phosphorylation leads to the recruitment of Shc and the activation of the MAPK/ERK signaling cascade (although there is cross-talk between the different pathways which is cell-specific)(23). The differential phosphorylation of these two tyrosines is believed to lead to different functional outcomes, and their differential phosphorylation in cells has already been used as an indicator of functional selectivity in EGFR signaling (16,25).

Experiments were performed with EGFR-mTurquoise (EGFR-mTurq), in which the fluorescent protein mTurq was attached to the C-terminus of EGFR via a 15 aa linker. This attachment does not impact the activation of EGFR (26). The cells were vesiculated and thousands of individual vesicles were imaged. To detect Y1068 phosphorylation, we used either anti-pY1068 or anti-pY1173 EGFR antibody, labeled with Alexa Fluor 488. The molar concentration of the antibodies always exceeded at least 5 times the total molar concentration of EGFR (~10 nM) and the pY-antibody dissociation constant (low nM). To start the reaction, we added ligands together with ATP kinase cocktail (1 mM ATP, 0.5 mM DTT, 10 mM MgCl_2_, 0.1 mM Na_3_VO_4_ (a phosphatase inhibitor)). The antibody was recruited to the vesicle membrane and the recruitment was quantified through the increase in membrane fluorescence using the neural network approach described in Supplemental Data. The imaging was performed at least 1 hour after the beginning of the reaction, based on kinetic traces of single vesicles which show complete equilibration after ~20 mins. Each vesicle was imaged using an automated microscope stage in two scans: one exciting mTurq at the C-terminus of EGFR, to assess EGFR concentration in each vesicle, and one exciting the fluorophore on the anti-pY antibody, to assess its concentration on the membrane in each vesicle (as it is a measure of the degree of phosphorylation).

Complete dose-response curves for WT EGFR Y1068 and Y1173 phosphorylation, in response to EGF, TGFα, and epiregulin, were collected (Figure 2A). A total of 12,225 vesicles were imaged, while the concentration of different ligands was varied from zero to saturating concentrations. For each individual vesicle, on the y axis we report the ratio of: (i) the fluorescence of AlexaF488, linked to the anti-pY antibody and (ii) the fluorescence of mTurq, linked to the receptor. Thus, on the y axis the values are proportional to the phosphorylated EGFR fraction, and on the x axis is the ligand concentration (Figure 2A). As all measurements utilized the same microscope setting and the same antibody batches, all Y1068 data are on same scale and all Y1173 data are on same scale.

**Figure 2.**
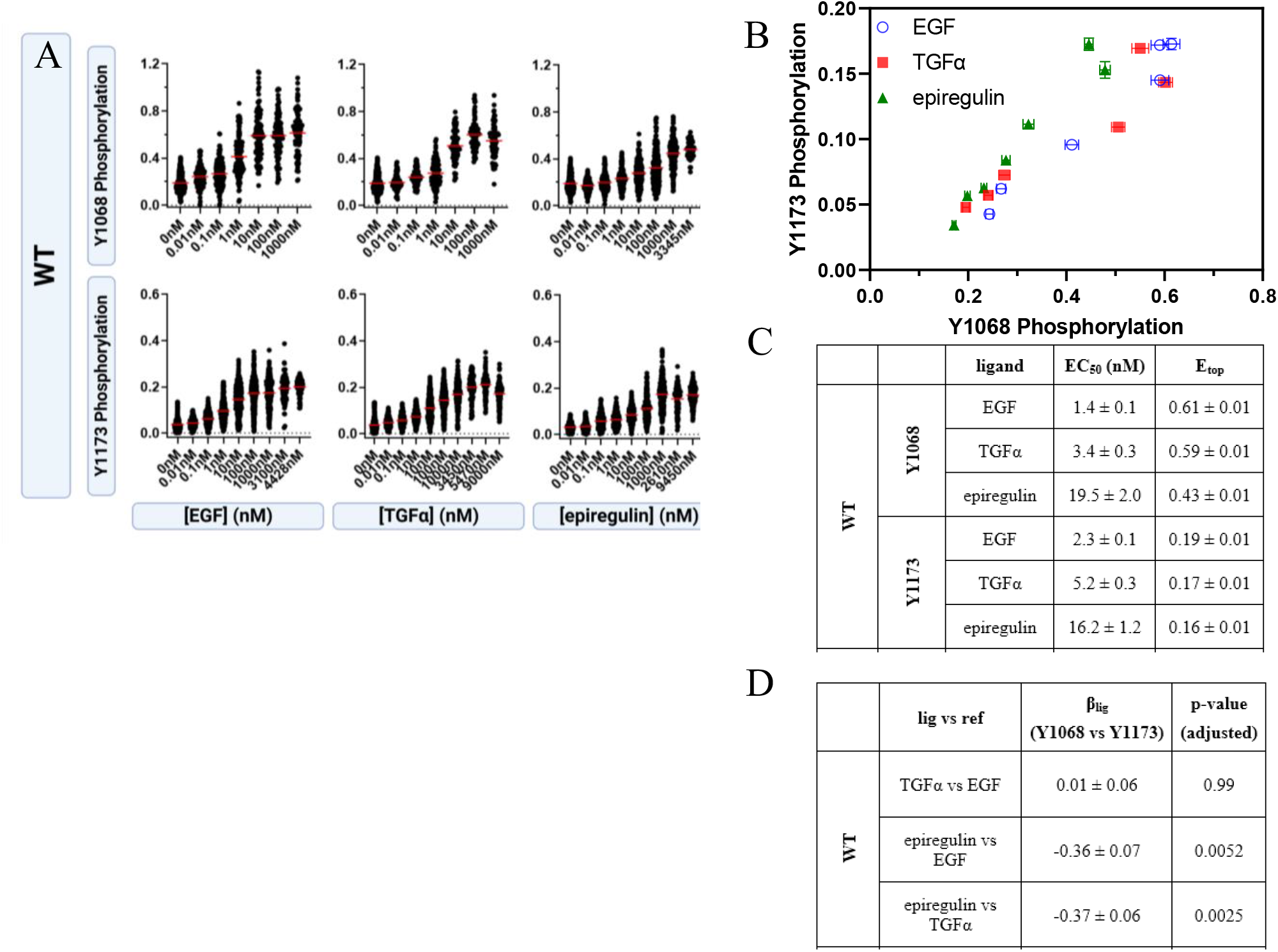
Ligand bias for WT EGFR. **(A)** Raw dose response curves for Y1068 and Y1173 phosphorylation, for the ligands EGF, TGFα, and epiregulin. Each point represents the ratio of either anti-pY1068 or anti-pY1173 fluorescence and EGFR-mTurq fluorescence for one individual vesicle. Each curve contains ~1,000 to ~3,000 data points (single vesicles). **(B)** Bias plots. Errors are standard errors (often smaller than symbols). The epiregulin points diverge from the EGF and TGFα points. **(C)** The fits to the corrected dose response curves in Figure S6 yield the potency (EC_50_) and efficacy (E_top_) for the three ligands. Errors are standard errors. **(D)** Bias coefficients and standard errors. Epiregulin is biased towards Y1173 phosphorylation, as compared to EGF and TGFα.

To determine if either Y1068 or Y1173 is preferentially phosphorylated by a ligand in comparison to another ligand, or whether there is no preference, we created bias plots (Figure 2B). In the bias plots one response (Y1173) is plotted against the second response (Y1068) at the same ligand concentrations. These bias plots report directly on the relative effectiveness of the ligands to produce the two responses, without the need for assumptions or mathematical modeling (12,27). The bias plots in Figure 2B appear different, indicating that there is bias. In particular, the epiregulin points diverge from the EGF and TGFα points, in the direction of Y1173 phosphorylation. Thus, the bias plot in Figure 2B demonstrates that epiregulin preferentially phosphorylates Y1173 over Y1068, when compared to EGF and TGFα.

Signaling bias due to a specific ligand can be identified and quantified with respect to a reference ligand by calculating bias coefficients (7, 15, 28, 29), such as the widely used β_lig_ given in equation (1) (7,12). This requires that we know the potencies, EC_50_, and the efficacies (maximum effects), E_top_, of the ligand and the reference ligand for the two responses, Y1068 and Y1173. The coefficient β_lig_ has a sign that indicates the preference of the ligand, as compared to the reference ligand, for a particular response (“+” if the first response is preferred and “–” if the second response is preferred), as well as a magnitude which reports on the degree of bias. The case of β_lig_ = 0 indicates that the ligand is not biased when compared to the reference ligand.

To calculate EC_50_ and E_top_ for the three ligands, we first note that the phosphorylation at zero ligand in Figure 2A is not zero. This is consistent with prior work, showing that EGFR can be phosphorylated in the absence of ligand, but cannot trigger downstream signaling (26), likely because the structure of the unliganded EGFR dimer is different from the structure of the active ligand-bound EGFR dimers (26). To characterize the response to ligand, we corrected the measured dose responses for the contribution of the unliganded dimers as described in the Supplement. To do so, we modeled the distributions of unliganded and liganded dimers based on the thermodynamic cycle shown in Figure S4. The measured single vesicle dose response curves are then fitted to equation (S18) to determine the potencies (EC_50_) and the efficacies (E_top_) of the ligands in inducing Y1068 and Y1173 phosphorylation. The values are given in Figures 2C and are used to calculate the bias coefficients in Figure 2D. A one-way ANOVA analysis of the bias coefficients in Figures 2D, followed by Tukey’s multiple comparison test, confirmed the presence of epiregulin-induced bias towards Y1173 phosphorylation, when compared to EGF (p = 0.005) and TGFα (p = 0.003). The corrected averaged dose response curves are shown in Figure S7.

### The common NSCLC L834R (L858R) driver mutation in EGFR induces intrinsic bias in signal propagation across the plasma membrane

NSCLC represents over 85% of all lung cancers and is associated with high mortality (30). The five-year survival for all stages of progression is less than 17%. This cancer is due to EGFR mutations in approximately 10–15% of Caucasian patients and in up to 50% of Asian patients. Of the single amino acid mutations, the L834R mutation is the most common one, accounting for about 40-45% of the cases where EGFR is mutated. (This mutation is often referred to as the “L858R mutation” when the EGFR signal peptide is counted).

We acquired dose response curves for L834R EGFR in response to EGF, TGFα, and epiregulin. A total of 7,923 individual vesicles were imaged and analyzed in these experiments (Figure 3A). We then constructed ligand bias plots (Figure 3B) and calculated bias coefficients (Figure 3D). In the case of L834R EGFR, both TGFα and epiregulin are biased towards Y1173 phosphorylation over Y1068 phosphorylation, as compared to EGF. These data demonstrate that the relative bias of the three EGFR ligands is altered due to the L834R mutation. The corrected averaged dose response curves are shown in Figure S7.

**Figure 3.**
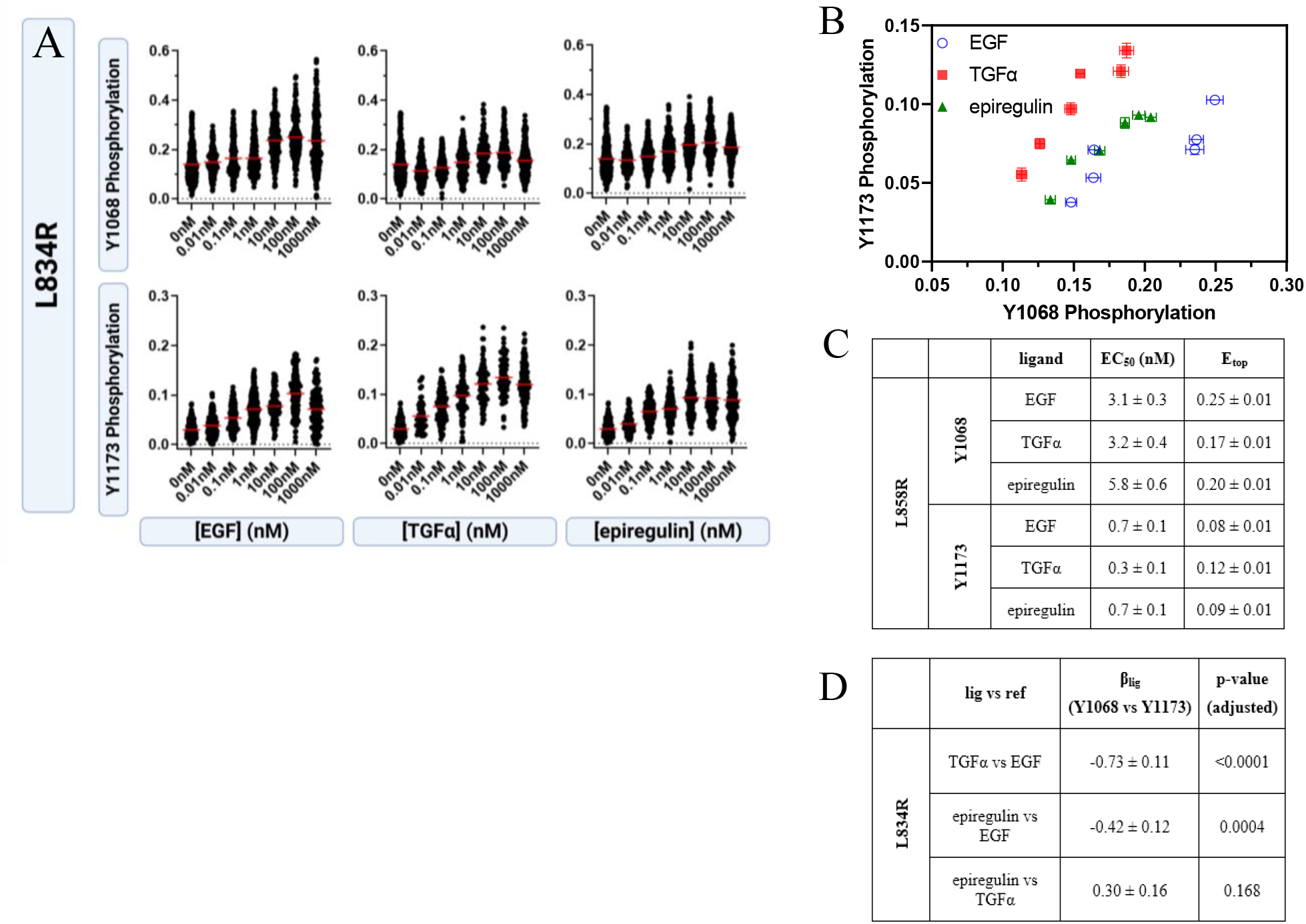
Ligand bias in L834R EGFR phosphorylation. **(A)** Single vesicle dose response curves for Y1068 and Y1173 phosphorylation, for EGF, TGFα, and epiregulin. Each point represents the ratio of either anti-pY1068 or anti-pY1173 fluorescence and EGFR-mTurq fluorescence for one individual vesicle. Each curve contains ~1,000 to ~1,800 data points. **(B)** Bias plots. Errors are standard errors (often smaller than symbols). **(C)** The fits to the dose response curves yield the potency (EC_50_) and efficacy (E_top_) for the three ligands. Errors are standard errors. **(D)** Bias coefficients and their standard errors.

To directly answer the question if the mutation causes bias in EGFR signaling, we created bias plots while directly comparing the wild-type and the mutant (Figure 4A-C). These are mutation-induced bias plots, distinctly different from the ligand bias plots in Figures 2B and 3B, as they now directly compare the mutant and the wild-type. In Figure 4A-C we see that the mutation induces significant preference for Y1173 phosphorylation over Y1068 phosphorylation, when compared to the wild-type, in the presence of the three ligands. We then calculated a novel type of bias coefficient, the mutation-induced bias coefficient β_mut_, to quantify the degree of bias introduced by the mutation in EGFR signaling in response to a specific ligand. The values were calculated using equation (2), where “response A” refers to Y1068 and “response B” refers to Y1173 phosphorylation (Figure 4D). The effect is largest in the case of TGFα, but highly statistically significant for all ligands, based on t-tests. This is a direct demonstration that the L834R mutation induces intrinsic bias in EGFR phosphorylation in the plasma membrane for all studied ligands. The corrected averaged dose response curves for the wild-type and the mutant are compared in Figure S8.

**Figure 4.**
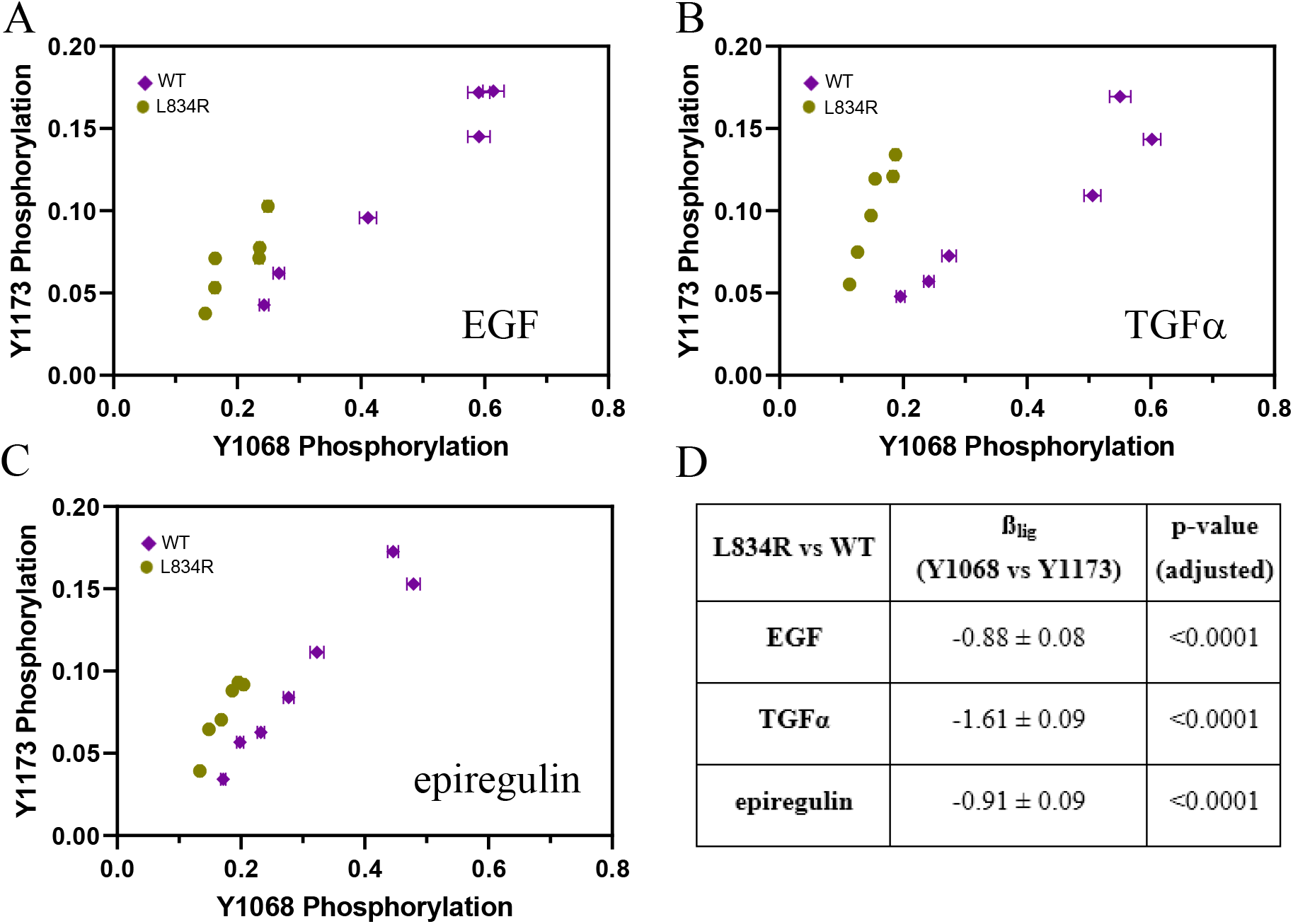
L834R mutation-induced bias. **(A)-(C)**. L834R-induced bias plots in the presence of EGF, TGFα, and epiregulin. Errors are standard errors (often smaller than symbols). **(D)** Bias coefficients and their standard errors. The L834R mutation induces statistically significant preference for Y1173 phosphorylation over Y1068 phosphorylation, as compared to the wild-type, in the presence of all three ligands.

### A measurement of the phosphorylation transducer function

The transducer function relates a response to the stimulus that is causing it (31). In our case, the response is the phosphorylation of a tyrosine in the intracellular domain of an RTK. The stimulus is the formation of the ligand-bound RTK dimers (31). We therefore sought to measure both ligand binding and phosphorylation simultaneously so we can plot one versus the other and obtain the transducer function.

To demonstrate the feasibility of transducer function measurements, we used commercially available EGF ligand from mouse that is labeled with rhodamine at its N-terminus (rho-mEGF, Thermofisher, E3481), a ligand that binds human EGFR with 3 times lower affinity than human EGF (32). To determine the transducer function for rho-mEGF, individual vesicles were imaged in a confocal microscope in three scans to measure: (i) the fluorescence of rhodamine, linked to mEGF, on the membrane, to quantify the bound ligand in the plasma membrane in each vesicle Ex:552nm; Em:565-625nm, (ii) the fluorescence of Alexa488, linked to the anti-phosphoY antibody, to quantify phosphorylated EGFR in the membrane Ex:488nm; Em:500-540nm (iii) the fluorescence of mTurq, linked to the receptor, in order to quantify EGFR in the plasma membrane in each vesicle Ex:448nm; Em:460-510nm. One vesicle, imaged in the three scans, in the presence of 5 nM EGF, is shown in Figure 5A. More than three thousand vesicles were imaged, while the ligand concentration was varied from zero to saturating concentrations.

**Figure 5.**
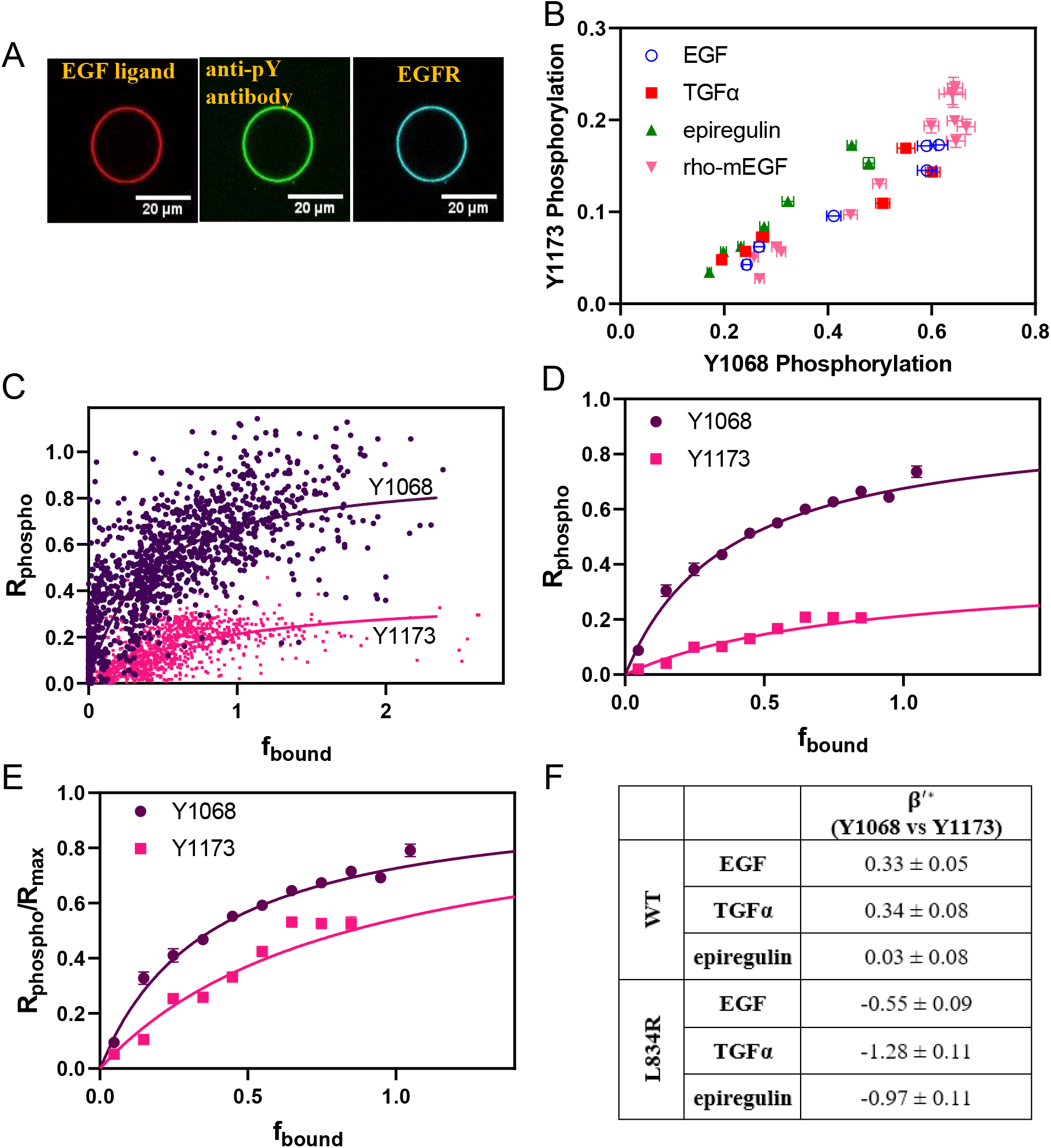
The rho-mEGF EGFR transducer function. **(A)** One vesicle imaged in three channels at 5 nM rho-mEGF in the presence of ATP/kinase cocktail. 3,085 individual vesicles were imaged and analyzed while the concentration of rho-mEGF was varied. **(B)** Bias plots for rho-mEGFR and human EGF, TGFα, and epiregulin. EGF and rho-mEGF are not biased ligands. **(C)** Phosphorylation response versus ligand-bound EGFR fraction for individual vesicles. The x axis is scaled such that the maximum average bound fraction is set to 1, and the y axis is phosphorylation corrected for constitutive (ligand-independent) phosphorylation. The solid lines are the transducer function fits (equation (S22)) to all the single vesicle data. **(D)** The single-vesicle data has been binned in an interval of 0.1 and is shown along with the fits to all the single vesicle data. Shown are standard errors; if not visible they are smaller than the symbols. **(E)** The normalized transducer function, given by equation (S24). **(F)** Absolute bias coefficients, calculated using equations (S36) to (S39).

The mouse and human EGF differ in sequence and affinity to human EGFR (See Supplemental Data). To assess if the mouse rho-EGF induces biased EGFR signaling, as compared to the three human ligands, we used the acquired phosphorylation dose response curves in Figure S9 to constructed bias plots comparing rho-mEGF and the human ligands (Figure 5B), and we calculated bias coefficients (Table S2). By ANOVA, the two EGF ligands are not biased, despite the reported differences in affinity to EGFR (32).

In Figure 5C, we plot the phosphorylation response as a function of the stimulus (ligand-bound EGFR fraction, f_bound_), for each individual vesicle that was imaged in the three channels. The x axis is scaled such that the maximum average fraction of ligand-bound EGFR is set to 1. This plot represents the transducer function.

We fit the data in Figure 5C using equation (S22) to determine (i) Rmax, the maximal possible signal that can be achieved in the experiment by a true full agonist (K_resp_→0), which depends on the fluorescent properties of the antibodies and (ii) K_resp_, the fraction of ligand-bound receptors that yields 50% of R_max_. The smaller the value of K_resp_, the more efficient the phosphorylation (see Supplemental Data). The best-fit values for Y1068 are K_resp_ = 0.40 ± 0.03 and R_max_ = 0.95 ± 0.03. The best-fit values for Y1173 are K_resp_ = 0.86 ± 0.08 and R_max_ = 0.39 ± 0.02. In Figure 5D, we show the fits along with the data, binned in intervals of 0.1 on the x axis. Only bins containing at least 50 vesicles are shown.

K_resp_ for Y1068 phosphorylation is the smaller of the two, indicating that Y1068 phosphorylation is more efficient than 1173 phosphorylation in response to EGF. Since the two R_max_ values differ because of the different fluorescent properties of the two antibodies, in Figure 5E we plot the normalized transducer function, i.e. the dependence of R_phospho_/R_max_ on the bound fraction, f_bound_. The y value at f_bound_=1 is the phosphorylation efficiency, calculated using equation (S23) as 0.71 ± 0.02 for Y1068 and 0.54 ± 0.02 for Y1173. Thus, rho-mEGF is a partial agonist for both responses. Human EGF, TGFα and epiregulin are also partial agonists, as the E_top_ values for their responses do not exceed the ones for rho-mEGF.

### Calculation of absolute bias coefficients

Bias coefficients calculated using equations (1) and (2) are relative, i.e. β_lig_ is always calculated with respect to the reference ligand in the litarature. However, the effective equilibrium constants K_resp_ can be used to calculate absolute bias coefficients, β’*_*lig*_, and β’*_*ref*_ (see equation (S38) and (S39)). First, we calculate β’*_rho-mEGF_ using equation (S37); β’*_rho-mEGF_ = 0.33 ± 0.50 Since this ligand is not biased in comparison to EGF, β’*_EGF_ = β’*_rho-mEGF_. The absolute β’*_EGF_ directly reports on the preference of a ligand towards either Y1068 or Y1173 phosphorylation. The value of β’*_EGF_ is positive, indicating that Y1068 is preferentially phosphorylated in response to EGF, as compared to Y1173.

With the values of β_lig_ and β’*_EGF_ known, we calculate the absolute bias coefficients β’*_TGFa_ and β’*_epiregulin_, for TGFα and epiregulin, using equation (S38). Similarly, we calculate the absolute bias coefficients β’*_mut_ for the L834R mutant using equation (S39). All absolute bias coefficients are shown in Figure 5F. We see that Y1068 in WT EGFR is preferentially phosphorylated in response to EGF and TGFα, but there is no preference in response to epiregulin. The signaling of the mutant is always biased towards Y1173 phosphorylation.

## DISCUSSION

Here we introduce a methodology to quantify (i) ligand and mutation-induced bias coefficients, both on relative and absolute scales and (ii) the transducer function describing RTK phosphorylation upon ligand stimulation. The critical descriptors of RTK activation measured here are “intrinsic”, as they describe the direct response of the RTKs to ligand binding, without contributions from downstream signaling feedback loops and system bias. This method can be used for all RTKs, and all membrane receptors in general.

The power of the methodology comes from the use of plasma derived vesicles produced via osmotic vesiculation. It has been proposed that defects in vesicles may form due to shear stress that ruptures the vesicle away from the cell membrane, or due to dysfunction of ESCRT components that mediate membrane scission (20). Protein dysfunction during scission could explain the much higher permeability of the osmotically-derived vesicles that we use as compared to the commonly used DTT/formaldehyde vesicles described in ref (20). Vesiculation occurs overnight in the osmotic vesiculation protocol, but takes only 2-3 hours in the DTT/formaldehyde protocol. After 12 hours of sustained stress, the activity of the ESCRT machinery could be significantly compromised.

The vesicles allow access of externally added antibodies to the phosphotyrosines on the RTK kinase domain, to allow for direct detection of phosphorylation. The vesicles lack cytoplasmic molecules involved in downstream signaling and thus there is no system bias in the measurements. The phosphorylation reaction is initiated by the researcher, by adding ligand and ATP kinase cocktail, and phosphorylation is followed through the recruitment of labeled specific anti-pY antibodies to the vesicle membranes. There is no signal attenuation because there is no RTK downregulation. Soluble phosphatases are not present, and the membrane phosphatases are inhibited, since the ATP kinase cocktail contains the inhibitors. Only mature RTKs in the plasma membrane are present. Antibodies, specific for only one tyrosine on only one RTK, verified in many RTK publications, are used in the detection. Data points in dose-response curves are derived from individual vesicles. Imaging is automated through the use of a commercial automated stage. Data processing is also automated using a neural network, and thus can be used for high-throughput inhibitor screening.

The data acquired with the new method can be compared to published data. First, epiregulin exhibits a lower potency for WT EGFR phosphorylation, as compared to EGF and TGFα (11, 33–36). Second, EGF and epiregulin are biased ligands, consistent with reports of ligand functional selectivity in cells (11). Thus, our results are consistent with data in the literature. As an important new development, the values of the intrinsic bias coefficients are now reported, and thus the degree of bias for multiple ligands is now quantified in the absence of feedback loops and system bias.

Another important result is the calculation of the EGF phosphorylation efficiency, which is the maximum possible phosphorylation that can be achieved in response to EGF. It is about 70% for Y1068 phosphorylation and 55% for Y1173 phosphorylation, Thus, EGF is not a full agonist, which suggests that new ligands can be designed to more strongly activate EGFR. This conclusion can be only drawn only after the efficiency of phosphorylation is measured, as done here.

We also gain insights into the origin of ligand bias in EGFR signaling. It has been argued that ligand bias in RTK signaling arises due to differential downregulation of the RTKs, or due to different abundances of cytoplasmic effectors (11,16). Here we show that bias arises in the first step of signaling transduction, along the length of the RTK.

We introduce a novel type of bias coefficient, β_mut_, which reports on mutation-induced bias, i.e. on the preferences of pathogenic RTK mutants to differentially phosphorylate tyrosines as compared to the wild-type RTKs. Mutation-induced bias is defined in analogy to ligand bias, where instead of comparing the effects of different ligands, we compare the wild-type and the mutant in the presence of the same ligand. By calculating both β_lig_ and β_mut_ from a comprehensive data set of dose-response curves, we uncouple and quantify biases introduced by ligand and by a pathogenic mutation. By simultaneously measuring the ligand binding and phosphorylation for a fluorescently labeled ligand, we quantify the characteristics of the transducer function, which ultimately allows the calculation of absolute intrinsic bias coefficients for natural ligands.

RTK mutations have been mainly classified as either gain of function (activating) or loss of function (deactivating) mutations (37). Here we show directly that the L834R EGFR mutation found in NSCLC induces bias in EGFR signal transduction across the plasma membrane. While EGFR signaling is biased towards Y1068 phosphorylation, the mutation switches the preference to Y1173 phosphorylation. It can be hypothesized that drug candidates that correct/unbias the first step in EGFR signal transduction can alter the signaling responses that are downstream from the mutant in a way that closely mimics WT EGFR signaling. We hope that the novel findings will create an impetus to quantify mutation-induced bias coefficients β_mut_ for the many known RTK pathogenic mutations, and to reclassify the mutations based on the sign and magnitude of the bias coefficients. This will pave the way for the development of novel mutation-specific inhibitors which account for the newly discovered complexity in RTK signaling.

## Materials and Methods

### Plasmid constructs

The plasmid encoding for human EGFR, tagged with the fluorescent protein mTurquoise (mTurq) at the C-terminus via a flexible GGS linker, is in the pSSX vector (38). The L834R mutation was introduced in EGFR using the QuikChange II Site-Directed Mutagenesis Kit according to manufacturer’s instructions (Agilent Technologies, #200523). The plasmid used for the neural network training encoded for the extracellular and transmembrane domain (ECTM) of FGFR, a (GGS)5 linker, and mTurq in the pcDNA3.1(+) vector (39). All plasmids were sequenced to confirm their identity (Genewiz).

### Cell culture and vesiculation

Chinese hamster ovary (CHO) cells were used in these experiments as they do not exhibit endogenous EGFR expression (40). CHO cells were cultured in Dulbecco’s modified Eagle medium (Gibco, #31600034) supplemented with 10% fetal bovine serum (HyClone, #SH30070.03), 1 mM nonessential amino acids, 10 mM D-glucose, and 18 mM sodium bicarbonate at 37 °C in a 5% CO_2_ environment. Cells were passed every other day using standard tissue culture techniques.

For vesiculation, the cells were seeded in a 6-well plate at a density of 2*10^4^ cells per well. 24 hours later, the cells were transfected with 1 or 1.5 μg plasmid DNA using FuGene HD (Promega, #E2311) according to the manufacturer’s protocol. 36 hours after transfection, vesiculation was induced using osmotic stress as described in (18). The osmotic pressure stresses the cells such that they release vesicles into solution without causing substantial cell detachment. The vesicles that were released in solution were collected by aspirating the supernatant with a cut 1000 μl micropipette tip.

### CHO vesicle characterization using dextran solutions

To characterize the permeability of CHO vesicles to macromolecules, FITC-labeled dextran was added to the vesicle solution and the ratio of FITC intensity inside and outside the vesicles was calculated for five different dextrans of sizes 20 – 2000 kDa. After vesiculation, CHO vesicles with ECTM FGFR3 tagged with mTurq incorporated in the plasma membrane were transferred to an 8-well glass bottom chamber slide (ibidi, #80827). 100 nM of FITC-labeled dextran in osmotic chloride salt buffer was added to the vesicles. The chamber slide was transferred to a TCS SP8 confocal microscope (Leica Biosystems, Wetzlar, Germany) equipped with an automated stage and a HyD hybrid detector in photon counting mode. The vesicles were allowed to settle for one hour. Image acquisition was automated by selecting pre-defined regions and focus points in the LAS X Navigator software (Leica Biosystems, Wetzlar, Germany). Two scans (256×256 pixels) per image were acquired, a ‘mTurq’-scan (λ=448 nm, emission window: 460 – 510 nm), where mTurq bound to FGFR3 is excited, and a ‘FITC’-scan (λ=488 nm, emission window: 500 – 540 nm), where FITC bound to dextran is excited. Images were acquired at 1% laser power with a 100 Hz scanning speed. To analyze the images, we developed a neural network approach as described in Supplementary Data.

### EGFR phosphorylation in CHO vesicles

To measure the phosphorylation of EGFR in the vesicle membranes, the vesicles were transferred to an 8-well glass bottom chamber slide (ibidi, #80827). The final concentration of the ATP cocktail ingredients were 1 nM ATP, 10 mM MgCl_2_, and 0.1 mM Na_3_VO_4_, a phosphatase inhibitor. The ligands used in this study were EGF (8916sf, Cell Signaling), TGFα (239A100, R&D Systems), Epiregulin (1195EP025 CF, R&D Systems), and EGF-tetramethylrhodamine (E3481, Thermofisher). For detection of Y1068 phosphorylation, 200 nM of Alexa488-labeled anti p-Y1068 EGFR antibody (IC3570G100, R&D Systems) was added. Y1173 phosphorylation was detected using 50 nM Alexa488-labeled anti phospho-Y1173 EGFR antibody (NBP1-44893AF488, Novus Biologicals).

Concentrations of the antibodies were chosen such that (i) the fluorescence intensities can be detected and measured on the plasma membrane (this depends on the labeling of the antibodies), (ii) the antibody amount exceeds the total amount of EGFR in the sample and (iii) the antibody concentration exceeds the anti-pY antibody dissociation constants (low nM). To determine the total EGFR concentration in a chamber slide well, 100 μl EGFR-mTurq vesicles were transferred to 96-well plates and full fluorescence emission spectra were collected with a H4 Synergy Hybrid Microplate Reader (BioTek Instruments, Winooski, VT). The samples were excited at 430 nm with a 9 nm bandwidth and the emitted fluorescence was collected from 450 – 620 nm with a 9 nm bandwidth with 5 nm steps. The emission spectra were corrected by subtracting the emission spectra of a vesicle sample derived from untransfected CHO cells. The maximum intensity of the corrected emission spectra at 475 nm was used to calculate the total EGFR concentration in a well upon calibration with purified solutions of mTurq of known concentration.

To monitor the reaction kinetics of EGFR phosphorylation, image acquisition was started right after the addition of the ligand/ATP cocktail to the vesicles. For dose response measurements, the phosphorylation reaction was allowed to reach equilibrium for 1 hour prior to image acquisition. Image acquisition was automated by selecting pre-defined regions and focus points in the LAS X Navigator software (Leica Biosystems, Wetzlar, Germany). The reaction was monitored for up to 5 hours. About 5000 images per experiment were acquired.

All images were acquired with a TCS SP8 confocal microscope (Leica Biosystems, Wetzlar, Germany) equipped with a motorized stage and a HyD hybrid detector in photon counting mode. Two scans per vesicle were taken, an ‘mTurq’-scan (λ=448 nm, emission window: 460 – 510 nm), where mTurq bound to EGFR is excited, and an ‘Alex488’-scan (λ=488 nm, emission window: 500 – 540 nm), where Alexa488 bound to the anti phospho antibody is excited. The images (512×512 pixels) were acquired at 1% laser power with a 50 Hz scanning speed.

### Ligand bias analysis

Dose response curves were fitted with the Hill equation with a slope of 1 (equation S17), as prescribed for calculations of the bias coefficient β_lig_ (7, 15, 28, 29). The best fit EC_50_ and E_top_ were used to calculate β_lig_ according to (7,12):

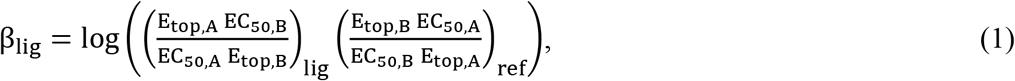

where response A is Y1068 phosphorylation and response B is Y1173 phosphorylation.

The mutation-induced bias coefficient was calculated as:

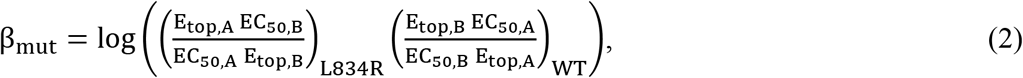

To test for ligand bias significance, a one way ANOVA followed by Tukey’s multiple comparisons test was performed using GraphPad Prism version 9.2.0 (GraphPad Software, San Diego, California USA).

The standard errors for bias coefficients and β’ values used in the statistical tests were derived from Monte-Carlo error estimations. For each parameter, 10^6^ normally distributed numbers were randomly generated using the mean and standard error of the parameter. The standard error of the distribution of the calculated bias coefficients was used for the statistical analysis.

## Acknowledgements

Supported by NIH R01GM068619 and NSF MCB 2106031. We thank Dr. M.D. Paul for helpful discussions and thermodynamic model development.

