## Supplementary Material for "Quantification of ligand and mutation-induced bias in EGFR phosphorylation in direct response to ligand binding"

### Identification of vesicles and quantification of their fluorescence intensities using machine learning

In this work, we had to analyze confocal images of thousands of vesicles. To increase the through-put of data processing, we developed an approach that uses artificial intelligence. The trained neural network is able to separate “vesicles” from “bad vesicles” and “cell debris”. A “vesicle” is defined as a spherical vesicle, completely separated from its parent cell (Figure 1C). Vesicles that are out of focus, cut off by image edges, non-circular, or attached to cells were classified as “bad vesicles”. Remnants of the cells after vesiculation were classified as “cell debris”.

For the neural network training, we used the imageLabeler app in Matlab to annotate rectangular regions in confocal micrographs for the classes (i) vesicle, (ii) bad vesicle, and (iii) cell debris. The resulting dataset of 1402 labeled objects was randomly shuffled and split into 70% training data and 30% test data. To build a neural network that identifies good vesicles, the FasterRCNN object detection network based on ResNet-18 was trained with the training data in Matlab. ResNet-18’s res4b_relu layer was chosen as the feature extraction layer. The learning rate was set to 0.001 and images were augmented during the training process by random horizontal reflection. 15 predefined anchor boxes, calculated using Matlab’s “estimateAnchorBoxes”, were used. Bounding box overlap ratios were set to 0.6 – 1 for positive training samples and to 0 – 0.3 for negative training samples to ensure a tight overlap with ground truth. The network was trained for a total of 10 epochs.

To train the network, we analyzed images of vesicles with FGFR3-eYFP, an RTK that has been used in prior work for method development (1, 2). Figure S1a shows the result of running the neural network on an image of FGFR3-eYFP vesicles. The neural network was able to distinguish between good and bad vesicles over broad intensity ranges. The receptor concentration was calculated for each detected vesicle as described (3) and the results are shown in Figure S1b. As a control, we labeled all vesicles in the test set by hand, and we compared the receptor concentration distribution to the one obtained with the computer vision-based code. The histogram of data in

Figure S1b from manual annotation shows good agreement with the histogram of the data from the neural network analysis. Thus, the neural network represents an efficient way to automatically analyze the images.

The neural network was used to quantify both EGFR phosphorylation in Figures 2A and 3A and ligand binding in Figure 5. It was also used to quantify the degree of penetration by dextrans into the vesicles in Figure 1B. The dextran experiments demonstrated the presence of defects of vesicles, which ensured that the phosphorylated tyrosines are accessible to externally added antibodies.

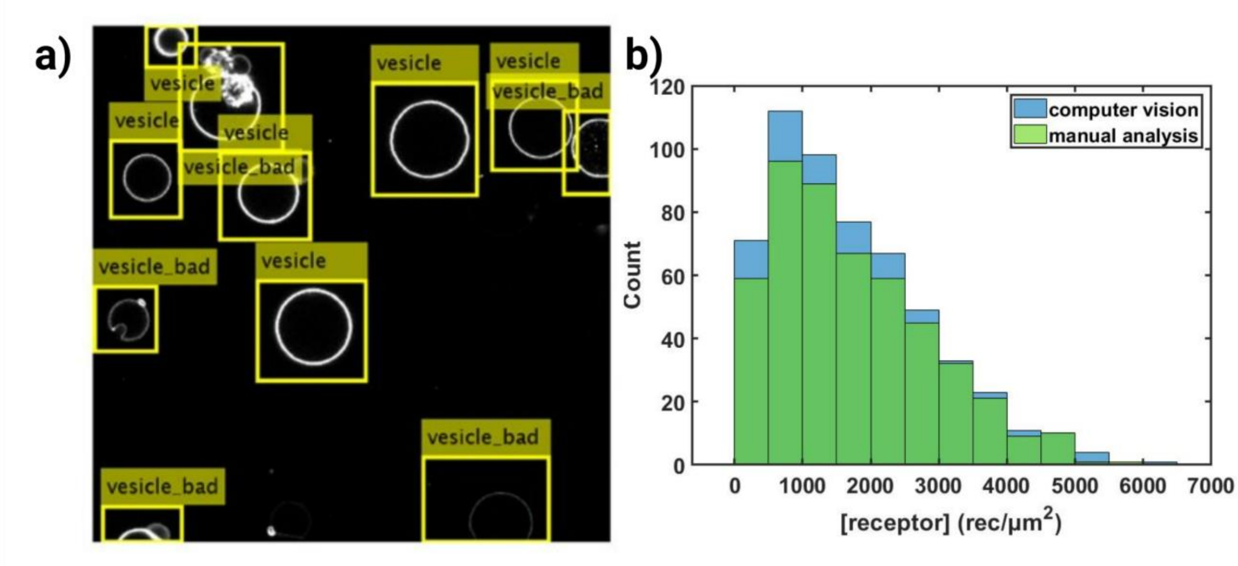


**Figure S1.** Neural network recognition of plasma-membrane derived vesicles with FGFR3-eYFP. a) Typical result of the analysis. The neural network is able to recognize good and bad vesicles of different intensities. b) Comparison of histograms of FGFR3-eYFP concentrations from vesicles detected by the computer-vision code (blue) and vesicles annotated manually (green). The two analysis methods show good agreement.

### Ligand bias

The term "ligand bias" describes the ability of ligands to preferentially activate a subset of signaling pathways, which can lead to fundamentally different biological responses (4, 5). Ligand bias does not simply reflect differences in potency ($EC_{50}$, the concentration of ligand that induces 50% of the maximum response), or efficacy ($E_{top}$, the maximum possible response due to a ligand). These differences represent quantitative differences in signaling, while bias represents fundamental (often called "qualitative") differences in signaling (6-8). This is illustrated in Figure S2 for two different ligands (blue and red) and two distinct responses (A, brown and B, green). Panels a and b show two cases of ligand bias while panels b and c show two cases of no bias. In Figure S2a both ligands activate response B similarly, but ligand 1 induces response A more efficiently than ligand 2. In Figure S2b ligand 1 preferentially induces response B while ligand 2 exhibits a preference towards response A. Figure S3b and c show only quantitative differences and therefore the ligands are not biased. Figure S3b depicts the case where response B is more efficiently triggered than response A, but the two ligands act similarly without bias. In Figure S3c ligand 1 activates both responses more efficiently than ligand 2, however the difference of activation in the two responses is the same for the two ligands and thus there is no bias. A variation of the last case would be where the efficiency of response B vary proportionally to the efficiency of response A. In such a case there would be no ligand bias, either.

The concept of ligand bias complements and expands concepts in traditional pharmacology where ligands are classified as either an agonist, antagonist or inverse agonist (9). A biased ligand can be either an agonist, antagonist or inverse agonist (10).

There have been numerous reports that propose the existence of ligand bias for FGF (11, 12), TRK (13-18), ERBB (19), Eph (20, 21), and PDGF (22) receptors. However, bias coefficients have not been calculated in these studies and the degree of the bias is unknown. Furthermore, it has been shown that different concentrations of adaptor proteins can bias signaling (23). This so-called "system bias" is often difficult to account for and decouple in cellular studies (24).

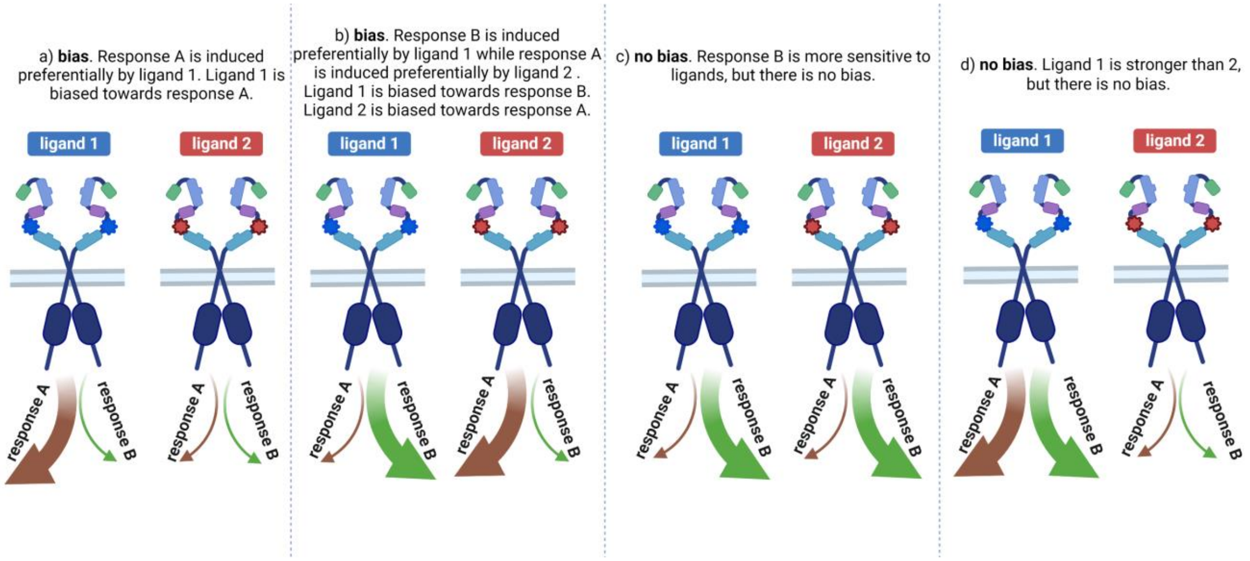


**Figure S2.** A schematic diagram showing two cases of bias (a and b) and two cases of no bias (c and d).

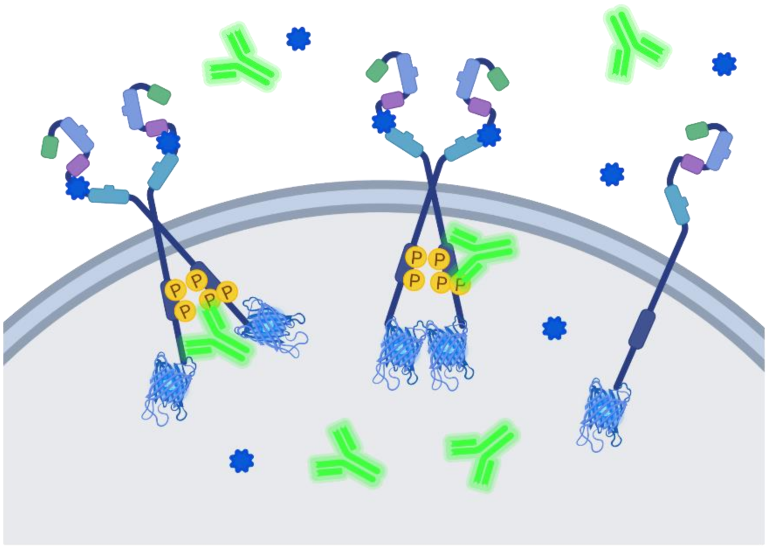

**Figure S3.** The experimental set up to follow receptor phosphorylation in plasma membrane derived vesicles which are permeant to macromolecules. RTKs in the vesicle membrane dimerize and get phosphorylated in response to ligand (blue star) binding. In the presence of ligand and ATP cocktail, a fluorescent specific anti-phosphoY antibody (green) is recruited to the membrane, and its recruitment is quantified, along with RTK concentrations in each vesicle.

### Correction of dose response curves for basal EGFR phosphorylation

According to the canonical model of RTK activation, RTKs are monomeric in the absence of ligand and are crosslinked upon ligand binding, which brings their catalytic domains in close proximity (26). However, numerous studies have demonstrated that EGFR activation is much more complex. First, it has been reported that EGFR can form dimers even in the absence of ligand, and that these dimers are phosphorylated, although they are not able to initiate downstream signaling cascades (27). Second, there are reports that EGFR can form oligomers upon ligand binding (28, 29). However, other studies show that EGFR exists predominantly in dimeric form (30).

To collect dose-response curves, phosphorylation of EGFR on the vesicle membrane was measured, as illustrated in Figure S3, over a broad range of ligand concentrations. These dose response curves, shown in Figures 2A and 3A in the main text, capture the phosphorylation of unliganded dimers, single-liganded dimers, and double-liganded EGFR dimers. The unliganded dimers account for basal phosphorylation of EGFR at zero ligand. Since we are interested in the response of EGFR to ligand, we modeled the concentration of unliganded EGFR dimers, and we corrected the dose response curves for the basal phosphorylation. This was done in order to calculate bias coefficients and therefore provide quantitative support for the bias plots, which rely on visual inspection (24).

Despite open questions about the association state of EGFR, it has been shown that its function can be described in quantitative terms by the thermodynamic cycle in Figure S4 which accounts for the coupling between ligand binding and EGFR dimerization (31). This cycle shows how increases in receptor and ligand concentration drive the formation of double-liganded EGFR dimers, and is used here to determine the unliganded dimer concentrations in each vesicle.

In our experiments, we know EGFR concentration in the membrane of each vesicle, $[R_t]$. We also know the total EGFR concentration in the imaging dish, $[R_{tot}]$, and the total ligand concentration in the imaging dish $[L_{tot}]$. The dimerization constants $K_D$ for EGFR and the L834R mutant have been measured as $K_D = 6.4 \times 10^{-3}$ μm$^2$ and $3.7 \times 10^{-2}$ μm$^2$, respectively (27). The EGF binding constants for the monomer $L_1$, the unliganded dimer $L_2$ and the liganded dimer $L_3$ have been measured as $L_1 = 4.6 \times 10^9$ M$^{-1}$, $L_2 = 5.3 \times 10^9$ M$^{-1}$, and $L_3 = 3.4 \times 10^8$ M$^{-1}$ (31). While both EGF and TGFα are known as "high-affinity ligands" for EGFR, with similar association constants, epiregulin is known as a low-affinity ligand, and its association constant has been reported to be 46 times lower than that of EGFR (19, 32). The mouse EGF-human EGFR association constant is 3 times lower than the human EGF association constant (33).

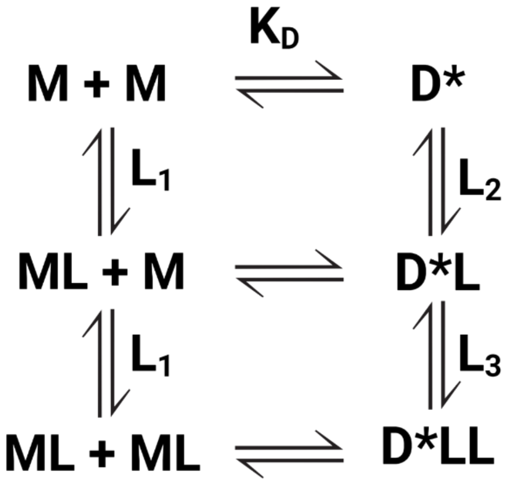


**Figure S4.** The thermodynamic cycle describing EGFR activation. * indicates that the dimeric receptors can be phosphorylated.

Based on the cycle in Figure S4, the concentrations of unliganded monomers, M, unliganded dimers, D, liganded monomers, ML, single-liganded dimers, DL, and double-liganded dimers, DLL, can be written as a function of these known equilibrium constants, the monomer concentration [M], and the free ligand concentration $[L_{free}]$ according to:

$$[D] = K_D * [M]^2 \tag{S1}$$

$$[ML] = L_1 * [L_{free}] * [M] \tag{S2}$$

$$[DL] = L_2 * [L_{free}] * K_D * [M]^2 \tag{S3}$$

$$[DLL] = L_2 * L_3 * [L_{free}]^2 * K_D * [M]^2 \tag{S4}$$

The total receptor concentration $[R_t]$ can be written as

$$[R_t] = [M] + [ML] + 2[D] + 2[DL] + 2[DLL] =$$
$$[M] + L_1 * [L_{free}] * [M] + 2K_D * [M]^2 + 2L_2 * [L_{free}] * K_D * [M]^2 + 2L_2 * L_3 * [L_{free}]^2 * K_D * [M]^2 (S5)$$

which can be rewritten as

$$[R_t] = (1 + L_1 * [L_{free}]) * [M] + (2K_D + 2L_2 * K_D * [L_{free}] + 2L_2 * L_3 * K_D * [L_{free}]^2) * [M]^2 (S6)$$

Thus, to be able to determine all the unknowns, we need to determine the free ligand concentration in the dish $[L_{free}]$ and the EGFR monomer concentration [M] in each vesicle.

First, we solve for the free ligand concentration in the dish. The total number of ligand molecules in the dish $L_{tot}$ is given by:

$$L_{tot} = [L_{free}] * volume + RL_{tot} \qquad (S7)$$

where $RL_{tot}$ the number of ligand molecules bound to receptors in the dish. "volume" is the volume of the vesicle solution in the dish.

$RL_{tot}$ is the sum of ligand molecules bound to monomers $ML_{tot}$, single-liganded dimers $DL_{tot}$, and double-liganded dimers $DLL_{tot}$:

$$RL_{tot} = ML_{tot} + DL_{tot} + 2DLL_{tot} \qquad (S8)$$

which can be written in terms of concentrations according to:

$$RL_{tot} = R_{tot} * \left(\frac{[ML_{tot}]}{[R_{tot}]} + \frac{[DL_{tot}]}{[R_{tot}]} + \frac{2[DLL_{tot}]}{[R_{tot}]}\right) \qquad (S9)$$

Next we assume that the average concentrations in the imaging dish is the same as the concentrations in a vesicle with an average total receptor concentration $[R_{avg}]$. We can write:

$$[R_{avg}] = \frac{\sum_{i=1}^{n} [M]_i + 2[D]_i + [ML]_i + 2[DL]_i + 2[DLL]_i}{n} \qquad (S10)$$

$$[ML_{avg}] = \frac{\sum_{i=1}^{n} [ML]_i}{n} \qquad (S11)$$

$$[DL_{avg}] = \frac{\sum_{i=1}^{n} [DL]_i}{n} \qquad (S12)$$

$$\left[DLL_{avg}\right] = \frac{\sum_{i=1}^{n}[DLL]_i}{n} \qquad (S13)$$

where $[x]_i$ is the concentration in the $i^{th}$ vesicle, and n the total number of vesicles. We can substitute (S11), (S12) and (S13) into $(S9)$ and write:

$$RL_{tot} = R_{tot} * \left(\frac{[ML_{avg}]}{[R_{avg}]} + \frac{[DL_{avg}]}{[R_{avg}]} + \frac{2[DLL_{avg}]}{[R_{avg}]}\right) \qquad (S14)$$

Note that in $(S14)$ the units of concentration are receptors per unit area instead of receptors per volume, as the receptors are confined in the 2-dimensional plasma membrane of the vesicle. Substituting equations (S1), (S2), (S3), and (S4) into $(S14)$, we obtain:

$$RL_{tot} = R_{tot} * \left(\frac{L_1 * [L_{free}] * [M_{avg}]}{[R_{avg}]} + \frac{2L_2 * [L_{free}] * K_D * [M_{avg}]^2}{[R_{avg}]} + \frac{4L_2 * L_3 * [L_{free}]^2 * K_D * [M_{avg}]^2}{[R_{avg}]}\right) \qquad (S15)$$

Substitution of (S15) into (S7) and further simplification yields:

$$L_{tot} = R_{tot} * \left(\frac{4L_2 * L_3 * K_D * [M_{avg}]^2}{[R_{avg}]}\right) * [L_{free}]^2 + \left(R_{tot} * \left(\frac{2L_2 * K_D * [M_{avg}]^2}{[R_{avg}]} + \frac{L_1 * [M_{avg}]}{[R_{avg}]}\right) + volume\right) * [L_{free}] \qquad (S16)$$

Equations $(S16)$ and $(S6)$ are two equations for the two unknowns, $[L_{free}]$ and $[M_{avg}]$. We use equation $(6)$, written for $[M_{avg}]$, to solve for $[M_{avg}]$ and we substitute it in equation $(S16)$. Then, we solve equation $(S16)$ for $[L_{free}]$ and we use this value to calculate [M] in each vesicle from equation (S6). With [M] and $[L_{free}]$ known, we calculate all other relevant concentrations using equations (S1) to (S4), for each vesicle.

The calculated concentrations, for every vesicle in the case of WT EGFR in the presence of EGF are plotted in Figure S8 as a function of the total ligand concentration and the total receptor concentration in the vesicles. At low ligand concentrations unliganded monomers and dimers dominate, while at high ligand concentrations mostly single-liganded monomers and double-liganded dimers are present. The presence of single-liganded dimers, DL, can only be observed at ligand concentrations around 10 nM EGF. The concentrations of unliganded dimers, [D], decreases with ligand concentration, as expected. Dividing each value of 2[D] by the total receptor concentration in the vesicle, [Rt], gives us the fraction of receptors in the unliganded dimeric state, $f_D$, in the vesicle:

$$f_D = \frac{2[D]}{[Rt]} \qquad (S17)$$

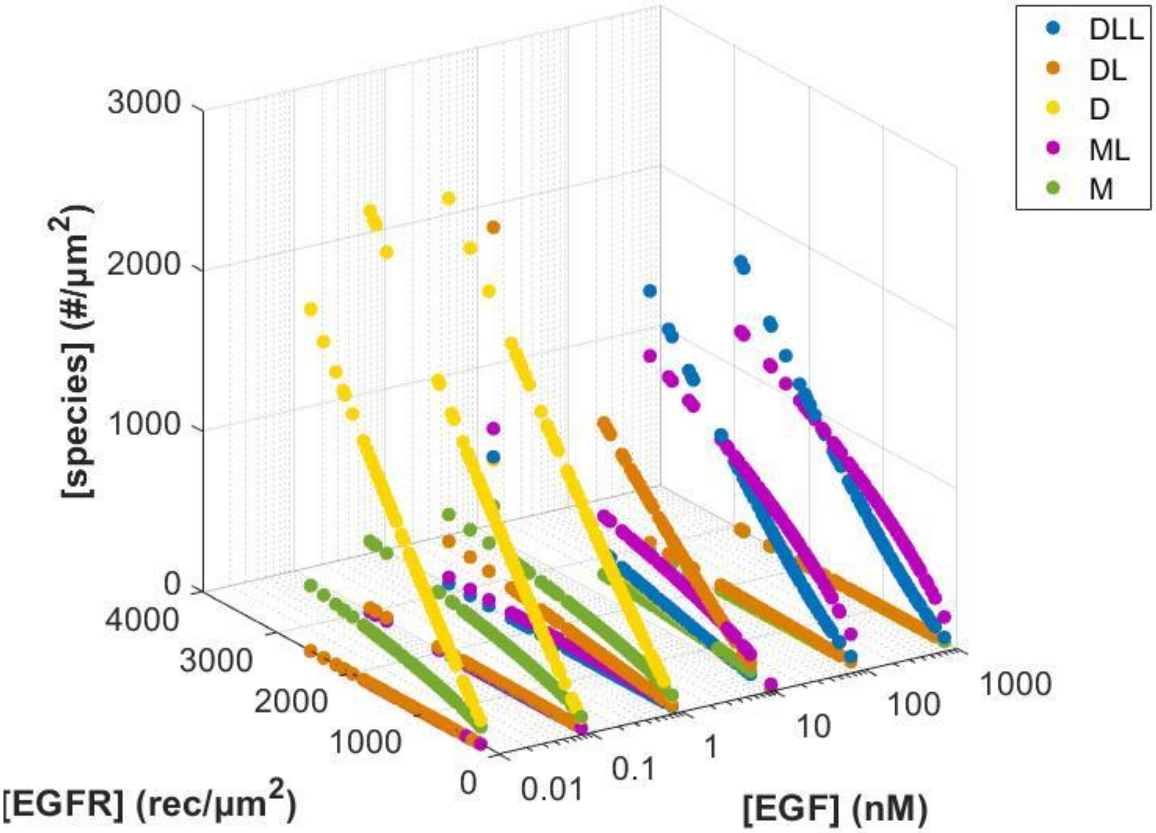


**Figure S5.** Concentrations of WT EGFR monomers and dimers, as a function of receptor and EGF concentrations. The monomer (green), unliganded dimer (yellow), liganded monomer (magenta), single-liganded dimer (orange), and double-liganded dimer (blue) distributions are plotted as a function of EGFR concentration (y-axis) and EGF concentration (x-axis) for every vesicle. The unliganded dimer concentrations (yellow) are used to correct the measured response in Figures 2A and 3A.

When no ligand is added to the vesicles, the observed phosphorylation, pY1068 and pY1173, is exclusively due to unliganded dimers. To quantify EGFR activation in direct response to ligand binding, $R_{phospho}$, we fit the data in Figure 2A and 3A in the main text to a Hill equation with n=1:

$$pY = R_{phosho} \ + k * f_D = \frac{[L_{tot}] * E_{top}}{EC_{50} + [L_{tot}]} + k * f_D \qquad (S18)$$

where $f_D$ is given by equation (S17) and

$$R_{phosho} = \frac{[L_{tot}] * E_{top}}{EC_{50} + [L_{tot}]} \quad (S19)$$

Here, pY is the measured phosphorylation, given by the fluorescence intensity of the anti-pY antibody signal divided by the EGFR-mTurq fluorescence, as shown in Figures 2A and 3A. $k$ is the best-fit correction factor, which is determined in the fit, along with $EC_{50}$ and $E_{top}$. The corrected dose response curves, $R_{phospho}$, are shown in Figure S6. The averaged dose response curves and the fits to all the single vesicle data are shown in Figures S7 and S8. The best-fit $EC_{50}$ and $E_{top}$ values are shown in Figures 2C and 3C.

The unliganded dimeric fraction, $f_D$ in equation (S18), depends on the ligand binding constants $L_1$, $L_2$, and $L_3$ (Figure S4). These constants have been measured for EGF (31). Here we use the same values for EGF and TGFα. Epiregulin is known as a low-affinity ligand, reported to bind 10 to 100 times weaker than EGF. By ITC, it has been shown to bind 46 times weaker (19). We use 46 times lower values for all three binding coefficients, L1, L2, and L3 for epiregulin-EGFR binding (19). For mouse EGF binding to EGFR, we use 3 times lower values for the three binding coefficients, L1, L2, and L3 (33).

.

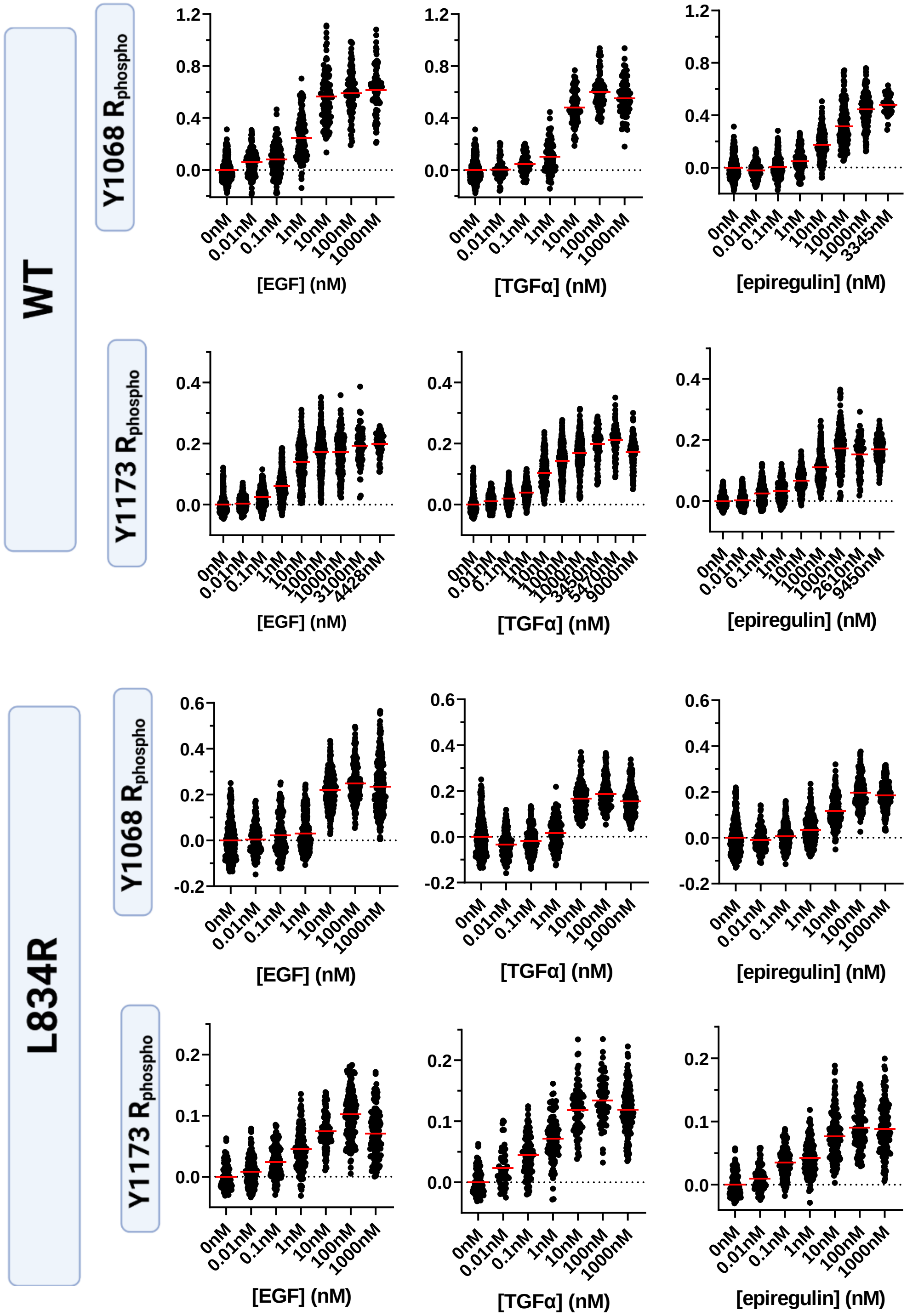


**Figure S6.** WT and L834R EGFR phosphorylation in individual vesicles in response to EGF, TGFα, and epiregulin, after correction for basal phosphorylation.

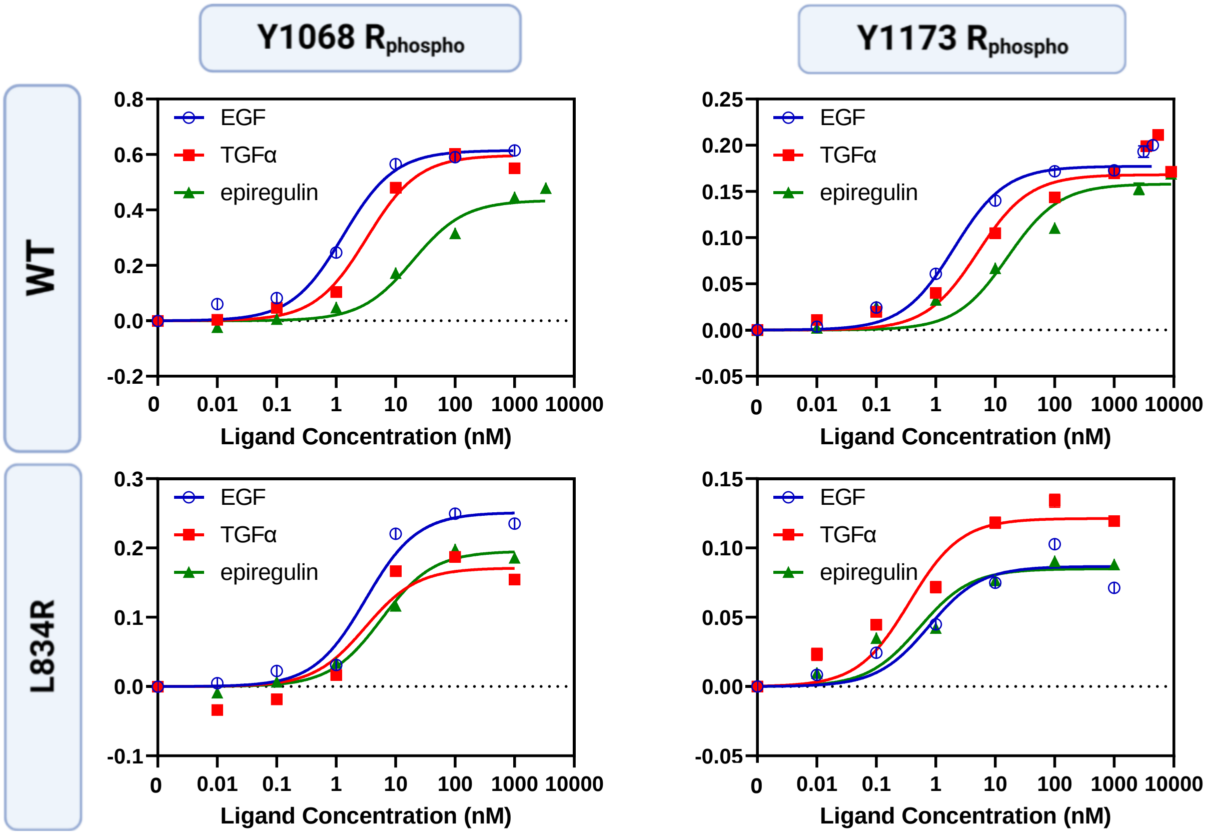


**Figure S7.** Fitted corrected dose-response curves for WT and L834R EGFR, for the three ligands. Shown are binned experimental data and standard errors (if errors are not visible, they are smaller than the symbols). The solid lines are fits to all the single vesicle data in Figures 2A and 3A.

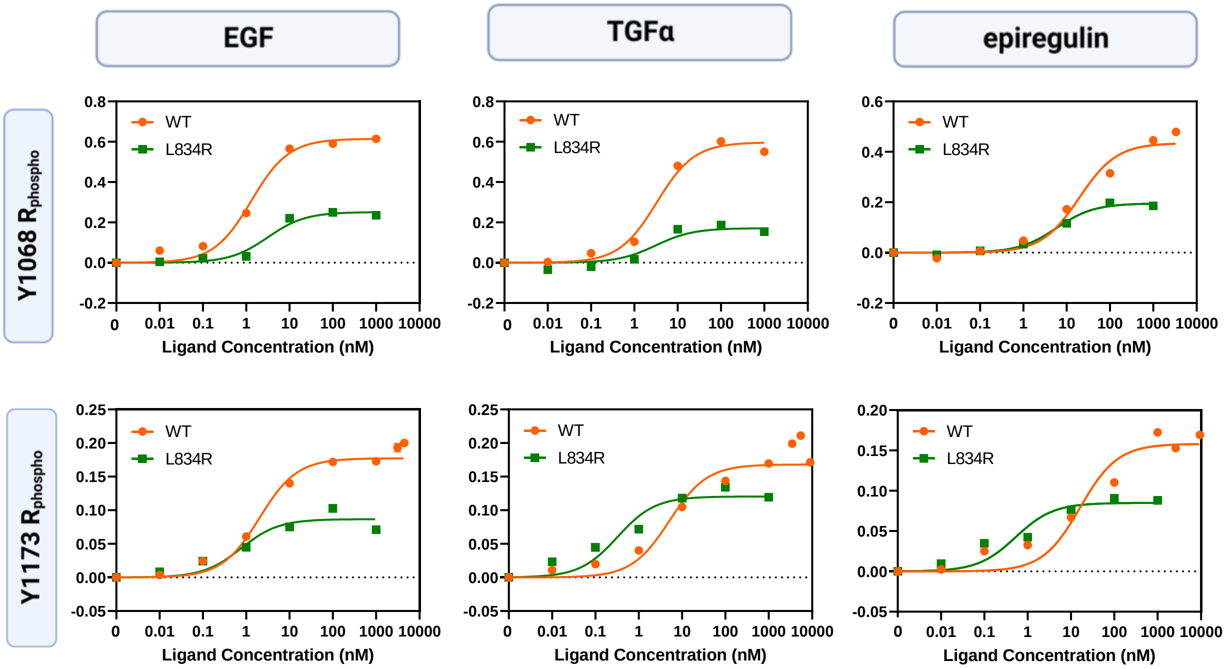


**Figure S8.** Comparison of corrected fitted dose response curves for WT and L834R EGFR in response to EGF, TGFα, and epiregulin. Same curves as in Figure S7, replotted to show the difference in phosphorylation between the wild-type and the mutant.

### The transducer function

The transducer function relates a response to the stimulus that is causing it. Experimentally, signaling responses downstream of a receptor depend on the abundance of activated receptors through a hyperbolic dependence (7, 34). The hyperbolic dependence was derived from first principles by Black and Leff (35), and is the basis for their "operational model", which is valid for different types of receptors including RTKs (25). In this model, the ligand-bound receptors act as a stimulus that activates the response with an effective equilibrium dissociation constant denoted as $K_{resp'}$ (6, 36):

$$\text{response} = \text{transducer function}(\text{stimulus}) = \frac{\text{stimulus}\ R_{max}}{\text{stimulus} + K_{\ resp'}} \qquad \text{(S20)}$$

In our experiments the "response" is the phosphorylation of a tyrosine in the intracellular domain of an RTK, $R_{phosho}$. We denote the maximum possible theoretical phosphorylation signal that can be achieved for this tyrosine in response to a full agonist as $R_{max}$ (37, 38).

The "stimulus" is the concentration of the ligand-bound receptors, [RL] (35). Therefore:

$$R_{phospho} = \frac{[RL]\ R_{max}}{[RL] + K_{resp'}} \qquad \text{(S21)}$$

If we divide both the numerator and denominator by the total receptor concentration, [Rt] and denote the fraction bound receptors, [RL]/[Rt], as $f_{bound}$, we obtain:

$$R_{phospho} = \frac{f_{bound}\ R_{max}}{f_{bound} + K_{resp}} \qquad \text{(S22)}$$

where

$$K_{resp} = \frac{K_{resp'}}{[Rt]} \quad \text{(S23)}$$

and the "stimulus" is now redefined as the fraction of ligand-bound receptors, $f_{bound}$. $K_{resp}$ is the fraction of ligand-bound receptors that yields 50% of $R_{max}$. The value of $R_{max}$ depends on the fluorescent properties of antibodies used for the detection and thus the phosphorylation response is fully described by the ratio of $R_{phospho}/R_{max}$:

$$\frac{R_{phospho}}{R_{max}} = \frac{f_{bound}}{f_{bound}+K_{resp}} \quad \text{(S24)}$$

The ligand-bound fraction $f_{bound}$ varies between 0 and 1. Setting $f_{bound}$=1, we define:

$$\text{phosphorylation efficiency} = \frac{R_{phospho}(f_{bound}=1)}{R_{max}} = \frac{1}{1+K_{resp}} \quad \text{(S25)}$$

This efficiency is the maximum phosphorylation fraction that can be achieved when all receptors are ligand-bound. It can be determined if the transducer function, given by equation (S22), is measured experimentally and $R_{max}$ and $K_{resp}$ are determined from a two-parameter fit. The smaller the value of $K_{resp}$, the more efficient the phosphorylation. Phosphorylation efficiency of $\rightarrow$ 1 ($K_{resp}$ $\rightarrow$ 0) is indicative of a full agonist.

#### The relation between $K_{resp}$ and bias, and the definition of absolute bias coefficients

Corrected dose-response curves are fitted using the Hill equation with n=1 (Eq. (S19)). The Black and Leff operational model is consistent with equation (S19), but also provides a physical-chemical description of the activation process (35). According to the Black and Leff model, the concentration of the ligand-bound receptor [RL] in equation (S21) depends on the concentrations of free receptor [R] and ligand [L], and on the effective ligand-receptor dissociation constant $K_L$ according to the equation:

$$[RL] = \frac{[R][L]}{K_L} \quad \text{(S26)}$$

The total receptor concentration [Rt] is:

$$[Rt] = [R] + [RL] \quad \text{(S27)}$$

Therefore:

$$[RL] = \frac{\frac{[Rt][L]}{K_L}}{1+\frac{[L]}{K_L}} \quad \text{(S28)}$$

Substitution of equation (S28) into equation (S21) yields:

$$\text{response} = \frac{[Rt][L]Rmax}{[Rt][L]+K_{resp}(K_L+[L])} = \frac{[Rt][L]Rmax}{[L]([Rt]+K_{resp})+(K_L K_{resp})} \quad \text{(S29)}$$

Dividing both the numerator and the denominator by $K_{resp}$, we obtain:

$$\text{response} = \frac{\left(\frac{[Rt]}{K_{resp}}\right)[L]Rmax}{\left(\frac{[Rt]}{K_{resp}}+1\right)[L]+(K_L)} = \frac{\tau[L]Rmax}{(\tau+1)[L]+(K_L)} \quad \text{(S30)}$$

where $\tau$ is the "transducer coefficient" defined as:

$$\tau = \frac{[\mathrm{Rt}]}{\mathrm{K}_{\mathrm{resp}}} \tag{S31}$$

Equation (S30) can also be written as:

$$\mathrm{response} = \frac{\tau/(\tau+1)[\mathrm{L}]\mathrm{Rmax}}{[\mathrm{L}]+(\mathrm{K_L})/(\tau+1)} \tag{S32}$$

Now we see that equation (S32) is the same as equation (S19), where

$$\mathrm{E_{top}} = \frac{\tau \mathrm{Rmax}}{(\tau+1)} \tag{S33}$$

$$\mathrm{EC_{50}} = \frac{\mathrm{K_L}}{(\tau+1)} \tag{S34}$$

We use equations (S33) and (S34) to arrive at an alternate expression of the bias coefficient

$$\beta_{\mathrm{lig}} = \log\left(\left(\frac{\mathrm{E_{top,A}\ EC_{50,B}}}{\mathrm{EC_{50,A}\ E_{top,B}}}\right)_{\mathrm{lig}}\left(\frac{\mathrm{E_{top,B}\ EC_{50,A}}}{\mathrm{EC_{50,B}\ E_{top,A}}}\right)_{\mathrm{ref}}\right) = \log\left(\left(\frac{\tau_A \mathrm{K_{L,B}}}{\mathrm{K_{L,A}}\tau_B}\right)_{\mathrm{lig}}\left(\frac{\tau_B \mathrm{K_{L,A}}}{\mathrm{K_{L,B}}\tau_A}\right)_{\mathrm{ref}}\right) \tag{S35}$$

Assuming that the ligand binding coefficient $\mathrm{K_L}$ does not depend on the binding of the anti-pY1068 and anti-pY1173 antibodies ($\mathrm{K_{L,A}= K_{L,B}}$), we arrive at

$$\beta_{\mathrm{lig}} = \log\left(\left(\frac{\tau_A}{\tau_B}\right)_{\mathrm{lig}}\left(\frac{\tau_B}{\tau_A}\right)_{\mathrm{ref}}\right) = \log\left(\left(\frac{\frac{[\mathrm{R_t}]}{\mathrm{K\ _{resp,A}}}}{\frac{[\mathrm{R_t}]}{\mathrm{K\ _{resp,B}}}}\right)_{\mathrm{lig}}\left(\frac{\frac{[\mathrm{R_t}]}{\mathrm{K\ _{resp,B}}}}{\frac{[\mathrm{R_t}]}{\mathrm{K\ _{resp,A}}}}\right)_{\mathrm{ref}}\right) =$$

$$\log\left(\left(\frac{\mathrm{K_{resp,B}}}{\mathrm{K_{resp,A}}}\right)_{\mathrm{lig}}\left(\frac{\mathrm{K_{resp,A}}}{\mathrm{K_{resp,B}}}\right)_{\mathrm{ref}}\right) = \quad +\log\left(\frac{\mathrm{K_{resp,B}}}{\mathrm{K_{resp,A}}}\right)_{\mathrm{lig}} - \log\left(\frac{\mathrm{K_{resp,A}}}{\mathrm{K_{resp,B}}}\right)_{\mathrm{ref}} = \beta'^{*}{}_{\mathrm{lig}} - \beta'^{*}{}_{\mathrm{ref}}$$

(S36)

where $\beta'^{*}{}_{\mathrm{lig}}$ and $\beta'^{*}{}_{\mathrm{ref}}$ are defined as:

$$\beta'^{*}{}_{\mathrm{lig}} = \log\left(\frac{\mathrm{E_{top,A}EC_{50,B}}}{\mathrm{EC_{50,A}E_{top,B}}}\right)_{\mathrm{lig}} = \log\left(\left(\frac{\mathrm{K_{resp,B}}}{\mathrm{K_{resp,A}}}\right)_{\mathrm{lig}}\right) = -\log\left(\left(\frac{\mathrm{K_{resp,A}}}{\mathrm{K_{resp,B}}}\right)_{\mathrm{lig}}\right) \quad \text{and}$$

$$\beta'^{*}{}_{\mathrm{ref}} = \log\left(\frac{\mathrm{E_{top,A}EC_{50,B}}}{\mathrm{EC_{50,A}E_{top,B}}}\right)_{\mathrm{ref}} = \log\left(\left(\frac{\mathrm{K_{resp,B}}}{\mathrm{K_{resp,A}}}\right)_{\mathrm{ref}}\right) = -\log\left(\left(\frac{\mathrm{K_{resp,A}}}{\mathrm{K_{resp,B}}}\right)_{\mathrm{ref}}\right) \tag{S37}$$

This definition of $\beta'^{*}{}_{\mathrm{lig}}$ and $\beta'^{*}{}_{\mathrm{ref}}$ does not include measurement bias and these coefficients report on the preference for phosphorylation in absolute terms. If $\mathrm{K_{resp,B}} > \mathrm{K_{resp,A}}$, then the value of $\beta'^{*}$ is positive and response A is preferred. If $\mathrm{K_{resp,A}} > \mathrm{K_{resp,B}}$, then the value of $\beta'^{*}$ is negative and response B is preferred.

Once $\beta'_{\mathrm{ref}}$ is known, we can calculate $\beta'^{*}{}_{\mathrm{lig}}$ as:

$$\beta'^{*}{}_{\mathrm{lig}} = \beta_{\mathrm{lig}} - \beta'^{*}{}_{\mathrm{ref}} \tag{S38}$$

By analogy, we can calculate the absolute bias coefficient $\beta'^{*}{}_{\mathrm{mut}}$ as:

$$\beta'^{*}{}_{\mathrm{mut}} = \beta_{\mathrm{mut}} - \beta'^{*}{}_{\mathrm{ref}}$$

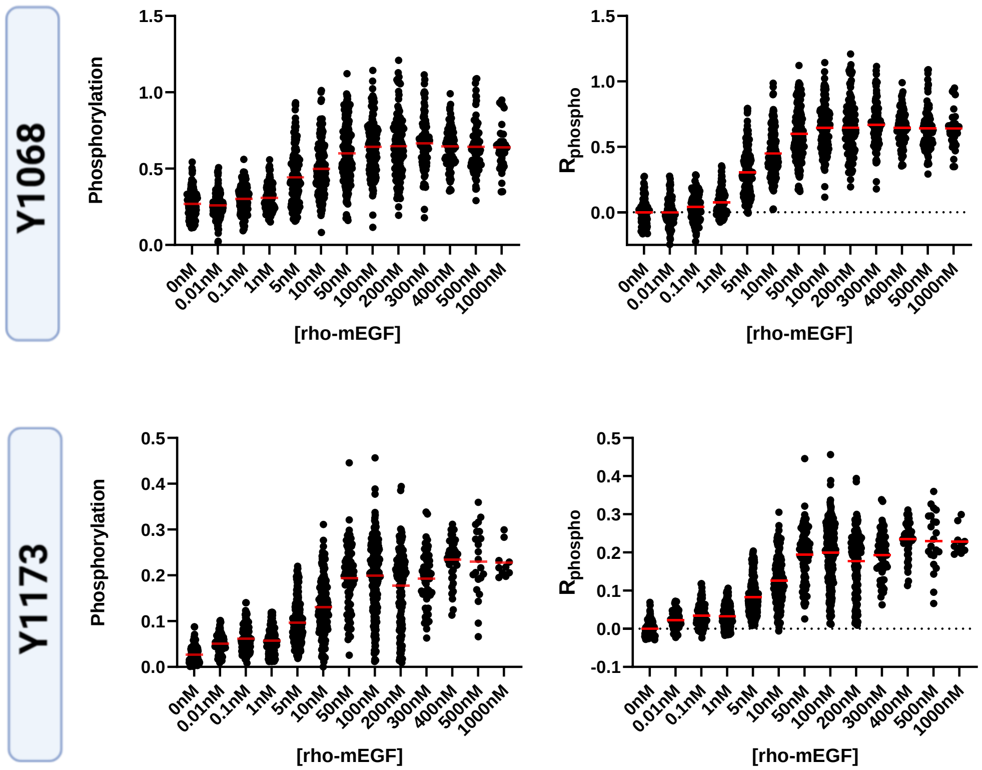


**Figure S9.** Measured and corrected EGFR dose responses to rho-mEGF.

| | **$EC_{50}$ (nM)** | **$E_{top}$** |
|---|---|---|
| **Y1068** | 5.5 ± 0.3 | 0.67 ± 0.01 |
| **Y1173** | 6.8 ± 0.4 | 0.21 ± 0.01 |

Table S1. $EC_{50}$ and $E_{top}$ for the rho-mEGF response.

| lig vs rho-mEGF | $ß_{lig}$ (Y1068 vs Y1173) | p-value (adjusted) |
|---|---|---|
| **EGF** | 0.12 ± 0.05 | 0.48 |
| **TGFα** | 0.13 ± 0.06 | 0.29 |
| **epiregulin** | -0.23 ± 0.06 | 0.0001 |

Table S2. Rho-mEGF bias coefficients.


1. Chen, L., J. Placone, L. Novicky, and K. Hristova. 2010. The extracellular domain of fibroblast growth factor receptor 3 inhibits ligand-independent dimerization. Science Signaling 3:ra86.
2. Sarabipour, S., R. B. Chan, B. Zhou, G. Di Paolo, and K. Hristova. 2015. Analytical characterization of plasma membrane-derived vesicles produced via osmotic and chemical vesiculation. Biochimica et Biophysica Acta 1848:1591-1598.
3. Chen, L. R., L. Novicky, M. Merzlyakov, T. Hristov, and K. Hristova. 2010. Measuring the Energetics of Membrane Protein Dimerization in Mammalian Membranes. Journal of the American Chemical Society 132:3628-3635.
4. Jarpe, M. B., C. Knall, F. M. Mitchell, A. M. Buhl, E. Duzic, and G. L. Johnson. 1998. [D-Arg1, D-Phe5, D-Trp7, 9, Leu11] Substance P acts as a biased agonist toward neuropeptide and chemokine receptors. Journal of Biological Chemistry 273:3097-3104.
5. Michel, M. C., and S. J. Charlton. 2018. Biased agonism in drug discovery—is it too soon to choose a path? Mol Pharmacol 93:259-265.
6. Kenakin, T. 2016. Measurement of Receptor Signaling Bias. Curr Protoc Pharmacol 74:2 15 11-12 15 15.
7. Kenakin, T. 2010. G protein coupled receptors as allosteric proteins and the role of allosteric modulators. J Recept Signal Transduct Res 30:313-321.
8. Kenakin, T., and A. Christopoulos. 2013. Signalling bias in new drug discovery: detection, quantification and therapeutic impact. Nat Rev Drug Discov 12:205-216.
9. Trevor, A. J. 2015. Chapter 2. Pharmacodynamics. In Katzung & Trevor's Pharmacology: Examination & Board Review, 11e. McGraw-Hill.
10. Berg, K. A., and W. P. Clarke. 2018. Making sense of pharmacology: inverse agonism and functional selectivity. International Journal of Neuropsychopharmacology 21:962-977.
11. Sarabipour, S., and K. Hristova. 2016. Mechanism of FGF receptor dimerization and activation. Nat Commun 7:10262.
12. Huang, Z., Y. Tan, J. Gu, Y. Liu, L. Song, J. Niu, L. Zhao, L. Srinivasan, Q. Lin, J. Deng, Y. Li, D. J. Conklin, T. A. Neubert, L. Cai, X. Li, and M. Mohammadi. 2017.

Uncoupling the Mitogenic and Metabolic Functions of FGF1 by Tuning FGF1-FGF Receptor Dimer Stability. Cell Rep 20:1717-1728.
13. Belliveau, D. J., I. Krivko, J. Kohn, C. Lachance, C. Pozniak, D. Rusakov, D. Kaplan, and F. D. Miller. 1997. NGF and neurotrophin-3 both activate TrkA on sympathetic neurons but differentially regulate survival and neuritogenesis. J Cell Biol 136:375-388.
14. Kuruvilla, R., L. S. Zweifel, N. O. Glebova, B. E. Lonze, G. Valdez, H. Ye, and D. D. Ginty. 2004. A neurotrophin signaling cascade coordinates sympathetic neuron development through differential control of TrkA trafficking and retrograde signaling. Cell 118:243-255.
15. Zaccaro, M. C., H. B. Lee, M. Pattarawarapan, Z. Xia, A. Caron, P.-J. L'Heureux, Y. Bengio, K. Burgess, and H. U. Saragovi. 2005. Selective small molecule peptidomimetic ligands of TrkC and TrkA receptors afford discrete or complete neurotrophic activities. Chem Biol 12:1015-1028.
16. Chen, D., F. Brahimi, Y. Angell, Y.-C. Li, J. Moscowicz, H. U. Saragovi, and K. Burgess. 2009. Bivalent peptidomimetic ligands of TrkC are biased agonists and selectively induce neuritogenesis or potentiate neurotrophin-3 trophic signals. ACS Chem Biol 4:769-781.
17. Harrington, A. W., C. St Hillaire, L. S. Zweifel, N. O. Glebova, P. Philippidou, S. Halegoua, and D. D. Ginty. 2011. Recruitment of actin modifiers to TrkA endosomes governs retrograde NGF signaling and survival. Cell 146:421-434.
18. Scarpi, D., D. Cirelli, C. Matrone, G. Castronovo, P. Rosini, E. G. Occhiato, F. Romano, L. Bartali, A. M. Clemente, G. Bottegoni, A. Cavalli, G. De Chiara, P. Bonini, P. Calissano, A. T. Palamara, E. Garaci, M. G. Torcia, A. Guarna, and F. Cozzolino. 2012. Low molecular weight, non-peptidic agonists of TrkA receptor with NGF-mimetic activity. Cell Death Dis 3:e339.
19. Freed, D. M., N. J. Bessman, A. Kiyatkin, E. Salazar-Cavazos, P. O. Byrne, J. O. Moore, C. C. Valley, K. M. Ferguson, D. J. Leahy, D. S. Lidke, and M. A. Lemmon. 2017. EGFR Ligands Differentially Stabilize Receptor Dimers to Specify Signaling Kinetics. Cell 171:683-695 e618.
20. Jorgensen, C., A. Sherman, G. I. Chen, A. Pasculescu, A. Poliakov, M. Hsiung, B. Larsen, D. G. Wilkinson, R. Linding, and T. Pawson. 2009. Cell-specific information processing in segregating populations of Eph receptor ephrin-expressing cells. Science 326:1502-1509.
21. Verheyen, T., T. Fang, D. Lindenhofer, Y. Wang, K. Akopyan, A. Lindqvist, B. Högberg, and A. I. Teixeira. 2020. Spatial organization-dependent EphA2 transcriptional responses revealed by ligand nanocalipers. Nucleic Acids Res 48:5777-5787.
22. Ho, C. C. M., A. Chhabra, P. Starkl, P. J. Schnorr, S. Wilmes, I. Moraga, H. S. Kwon, N. Gaudenzio, R. Sibilano, T. S. Wehrman, M. Gakovic, J. T. Sockolosky, M. R. Tiffany, A. M. Ring, J. Piehler, I. L. Weissman, S. J. Galli, J. A. Shizuru, and K. C. Garcia. 2017. Decoupling the Functional Pleiotropy of Stem Cell Factor by Tuning c-Kit Signaling. Cell 168:1041-1052 e1018.
23. Salazar-Cavazos, E., C. F. Nitta, E. D. Mitra, B. S. Wilson, K. A. Lidke, W. S. Hlavacek, and D. S. Lidke. 2020. Multisite EGFR phosphorylation is regulated by adaptor protein abundances and dimer lifetimes. Mol Biol Cell 31:695-708.
24. Kolb, P., T. Kenakin, S. P. H. Alexander, M. Bermudez, L. M. Bohn, C. S. Breinholt, M. Bouvier, S. J. Hill, E. Kostenis, K. A. Martemyanov, R. R. Neubig, H. O. Onaran, S.

Rajagopal, B. L. Roth, J. Selent, A. K. Shukla, M. E. Sommer, and D. E. Gloriam. 2022. Community guidelines for GPCR ligand bias: IUPHAR review 32. Br J Pharmacol 179:3651-3674.
25. Karl, K., M. D. Paul, E. B. Pasquale, and K. Hristova. 2020. Ligand bias in receptor tyrosine kinase signaling. J Biol Chem.
26. Weiss, A., and J. Schlessinger. 1998. Switching signals on or off by receptor dimerization. Cell 94:277-280.
27. Byrne, P. O., K. Hristova, and D. J. Leahy. 2020. EGFR forms ligand-independent oligomers that are distinct from the active state. The Journal of biological chemistry.
28. Needham, S. R., S. K. Roberts, A. Arkhipov, V. P. Mysore, C. J. Tynan, L. C. Zanetti-Domingues, E. T. Kim, V. Losasso, D. Korovesis, M. Hirsch, D. J. Rolfe, D. T. Clarke, M. D. Winn, A. Lajevardipour, A. H. Clayton, L. J. Pike, M. Perani, P. J. Parker, Y. Shan, D. E. Shaw, and M. L. Martin-Fernandez. 2016. EGFR oligomerization organizes kinase-active dimers into competent signalling platforms. Nat Commun 7:13307.
29. Huang, Y., S. Bharill, D. Karandur, S. M. Peterson, M. Marita, X. Shi, M. J. Kaliszewski, A. W. Smith, E. Y. Isacoff, and J. Kuriyan. 2016. Molecular basis for multimerization in the activation of the epidermal growth factor receptor. Elife 5.
30. Chung, I., R. Akita, R. Vandlen, D. Toomre, J. Schlessinger, and I. Mellman. 2010. Spatial control of EGF receptor activation by reversible dimerization on living cells. Nature 464:783-U163.
31. Macdonald, J. L., and L. J. Pike. 2008. Heterogeneity in EGF-binding affinities arises from negative cooperativity in an aggregating system. Proceedings of the National Academy of Sciences of the United States of America 105:112-117.
32. Sanders, J. M., M. E. Wampole, M. L. Thakur, and E. Wickstrom. 2013. Molecular determinants of epidermal growth factor binding: a molecular dynamics study. PLoS One 8:e54136.
33. Nexo, E., and H. F. Hansen. 1985. Binding of epidermal growth factor from man, rat and mouse to the human epidermal growth factor receptor. Biochim Biophys Acta 843:101-106.
34. Kenakin, T. 2017. Signaling bias in drug discovery. Expert Opin Drug Discov 12:321-333.
35. Black, J. W., and P. Leff. 1983. Operational models of pharmacological agonism. Proc R Soc Lond B Biol Sci 220:141-162.
36. Gundry, J., R. Glenn, P. Alagesan, and S. Rajagopal. 2017. A Practical Guide to Approaching Biased Agonism at G Protein Coupled Receptors. Front Neurosci 11:17.
37. Ehlert, F. J. 2018. Analysis of Biased Agonism. Prog Mol Biol Transl Sci 160:63-104.
38. Kenakin, T. 2017. A Scale of Agonism and Allosteric Modulation for Assessment of Selectivity, Bias, and Receptor Mutation. Mol Pharmacol 92:414-424.
